# Regulation of eIF6 through multisite and sequential phosphorylation by GSK3β

**DOI:** 10.1101/476978

**Authors:** Courtney F. Jungers, Jonah M. Elliff, Daniela S. Masson-Meyers, Christopher J. Phiel, Sofia Origanti

## Abstract

Eukaryotic translation initiation factor 6 is essential for the synthesis of 60S ribosomal subunits and for regulating the association of 60S and 40S subunits. A mechanistic understanding of how eIF6 modulates translation in response to stress, specifically starvation-induced stress, is lacking. Our studies have uncovered a novel mode of eIF6 regulation by GSK3 that is predominantly active in response to serum starvation. Human eIF6 is phosphorylated by GSK3β at a multisite motif in the C-terminus. Sequential phosphorylation by GSK3β requires phosphorylation at a priming site. In response to serum starvation, eIF6 accumulates in the cytoplasm and this altered localization is dependent on GSK3. Disruption of eIF6 phosphorylation enhances translation inhibition and markedly sensitizes the cells to serum starvation. These results suggest that eIF6 regulation by GSK3β contributes to the attenuation of global protein synthesis that is critical for adaptation to starvation-induced stress.

## Introduction

Eukaryotic initiation factor 6 (eIF6) is a key modulator of translation initiation that regulates the biogenesis and availability of the 60S ribosomal subunits^1–3^. eIF6 directly associates with pre-60S complexes in the nucleolus and is exported into the cytoplasm in complex with the 60S where it aids in 60S maturation^4–8^. The well-characterized role of eIF6 is its anti-association activity that prevents interactions between the 60S and 40S ribosomal subunits^2,9–12^. Structural and biochemical studies indicate that eIF6 binds to 60S at the 40S-binding interface and sterically hinders association of the 40S subunits^10–16^. Release of eIF6 from the mature 60S allows association of the 40S-mRNA complex, which leads to active 80S formation and translation initiation^10–15^. An inhibition of eIF6 release (or premature release) from the 60S subunits can greatly influence intersubunit association and translation initiation. A block in eIF6 release leads to an increase in the fraction of eIF6-bound 60S subunits that are unable to join 40S, which hinders the assembly of active 80S monosomes^10–13,15,16^. Alternatively, insufficient levels of eIF6 lead to spurious association of 60S with the 40S subunits that are devoid of mRNA. This causes an increase in the assembly of inactive 80S monosomes, which impairs translational response to growth stimuli as seen in *eIF6*+/- mice^17–19^. Thus, an impairment of eIF6 function can substantially affect ribosome levels, limit protein synthesis specifically in response to growth stimuli, and this deregulation has been shown to contribute to the underlying pathologies of diseases such as Shwachman-Bodian-Diamond syndrome, cancer and certain metabolic disorders^13,17–20^.

In response to stimuli such as insulin or phorbol esters, primary hepatocytes and mouse embryonic fibroblasts (MEFs) derived from the *eIF6*+/- mice do not upregulate protein synthesis unlike the wild type (WT) cells^17–19^. Similar defects in translation are also observed *in vivo,* where livers of *eIF6*+/- mice are smaller, and exhibit an accumulation of inactive/empty (mRNA-free) 80S ribosomes compared to WT^17–19^. The lack of insulin-dependent stimulation of protein synthesis in the *eIF6*+/- mice has been associated with a reprogramming of fatty acid synthesis and glycolytic pathways with implications for muscle, liver and fat metabolism^19,21^.

In terms of mechanism, the increased formation of inactive 80S complexes in the *eIF6*+/- cells is attributed to an impairment of its anti-association function in the cytoplasm^17–19^. Interestingly, such an accumulation of inactive 80S monosomes is commonly observed in cells subjected to stress, especially stress induced by nutrient deprivation or limitation^22–24^. Starvation or nutrient limitation in yeast and mammalian cells invokes an adaptive metabolic response that conserves energy by restricting global protein synthesis and leads to an accumulation of inactive (empty) 80S ribosomes along with an increase in the pool of free 60S subunits ^22–33^. The role of eIF6 as a ribosome anti-association factor and 60S biogenesis factor in modulating this starvation response has not been thoroughly explored. Given the key role of eIF6 in regulating translational response to insulin and growth factors in an mTOR-independent manner^19,34^, it is important to understand the mechanisms that control eIF6 in response to growth inhibitory stress responses. Towards addressing this, here we report a novel regulation of eIF6 by the Glycogen Synthase Kinase-3 (GSK3) that is active under conditions of serum starvation-induced stress.

Global proteomic and biochemical studies indicate that eIF6 is phosphorylated at multiple sites and majority of these sites are conserved and cluster around the C-terminal tail^35–41^. However, most of these phosphorylation sites have not been validated *in vivo* and the identity of the associated kinases have not been clarified. It is also unclear if these uncharacterized phosphorylation sites carry any functional relevance. Therefore, we carried out a sequence analysis and identified a putative motif for GSK3 phosphorylation within the C-terminal tail of eIF6. Previous studies show that GSK3 is uniquely activated in response to growth inhibitory conditions such as starvation or quiescence and is inhibited in response to insulin and other growth stimulatory conditions by AKT and mTORC1/p70S6K1-dependent phosphorylation^42–48^. GSK3 plays a prominent role in translational control by inhibiting the nucleotide exchange function of the translation initiation factor: eIF2B, which is reversed in response to insulin^42,49–51^. GSK3 phosphorylates the Ser-540 site in the epsilon subunit of eIF2B to inhibit its activity^42,49–51^. Here, we show that GSK3 also regulates eIF6 by phosphorylating human eIF6 on a multisite motif. GSK3 interacts with endogenous eIF6 *in vivo* and phosphorylation by GSK3β alters the subcellular localization of eIF6 in response to serum starvation. Interestingly, the phosphodead mutant cells exhibit increased levels of free 60S subunits and inactive 80S monosomes relative to the WT. Altering the levels of ribosomal subunits can have detrimental effects as seen in several ribosomopathies^52,53^. Here we show that the phosphodead mutant exhibit an enhanced inhibition of translation and are much more sensitive to serum starvation than WT. Based on these findings that eIF6 is a novel substrate of GSK3, we propose a model wherein GSK3-dependent phosphorylation of eIF6 contributes to translation inhibition in response to serum starvation by regulating 60S availability.

## Results

### The C-terminal tail of eIF6 is phosphorylated by GKS3β

Sequence analysis of the amino acid residues in the C-terminal tail of human eIF6 revealed the presence of sequential S/TXXXS/T motifs as potential candidates for phosphorylation by GSK3. We identified two such multisite motifs (motif 1 and 2) in the C-terminus that are highly conserved in higher eukaryotes from *Xenopus* to mammals (Fig. 1A) but show variability in lower eukaryotes such as yeast. GSK3 phosphorylates residues on its substrates in a sequential manner starting with the C-terminus, and for a majority of substrates, GSK3 works in concert with a priming kinase that phosphorylates a priming site located 4 residues away from the GSK3 recognition site (Fig. 1A)^42,44,45,54–56^. The potential priming sites are indicated in Fig. 1A. To determine if eIF6 is a substrate of GSK3, we immunoprecipitated eIF6 from HCT116 cells that were briefly serum starved for 4 hours to ensure that GSK3 and any associated priming kinase were activated (Fig. S1A, B). We confirmed that GSK3β was activated in response to short-term serum starvation in HCT116 cells by probing for the loss of the inhibitory phosphorylation of the serine-9 residue. The serine-9 site on GSK3β is phosphorylated by kinases such as AKT, and the phosphorylated N-terminal end acts as a pseudo substrate to inhibit GKS3β activity^43,45–48,54,55^. Therefore, a loss of S9 phosphorylation indicates activation of GSK3β as seen during serum starvation. At 4 hours of serum starvation, we observed that only a fraction of GSK3 is activated, indicating that at this time point the sites on eIF6 would not be fully occupied (Fig. S1A, B). Also, we did not use recombinant eIF6 purified from *E. coli* as the pure protein would not be in a primed state, and it was unclear if priming of eIF6 was a requisite for phosphorylation by GSK3. We also focused on the GSK3β isoform in this study, which is the well characterized isoform^43^.

**Figure 1.**
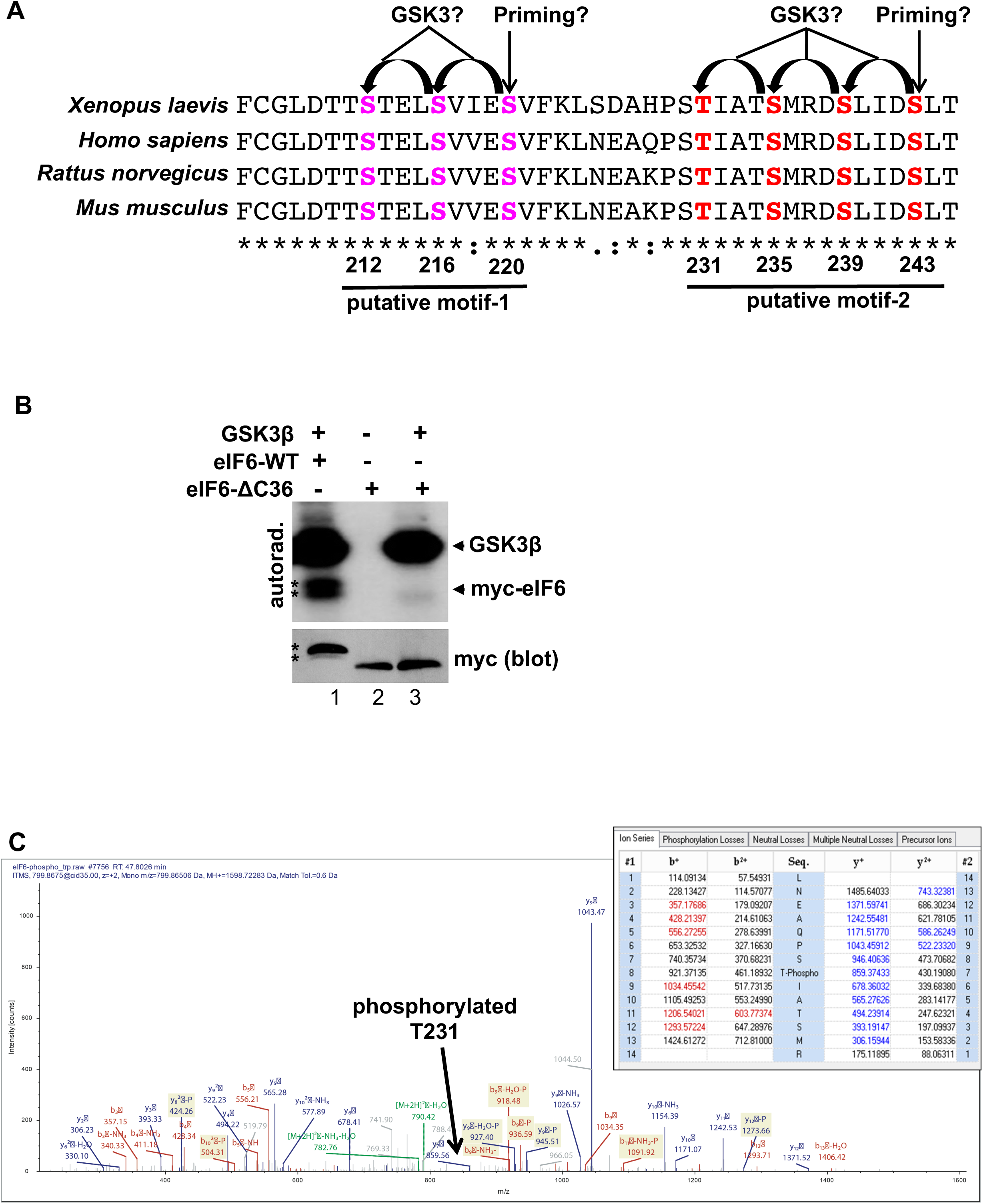
GSK3β phosphorylates a multisite motif in the C-terminal tail of eIF6. **A.** Sequence alignment of the last 41 amino acid residues (205 to 245) in the C-terminal tail of human eIF6 (NP_001254739) with *Xenopus* (NP_001083080.1), mouse (NP_034709.1), and rat eIF6 (NP_001032429). The conserved residues are marked with an asterix. Putative GSK3-specific phospho sites and priming site in multisite motif-1 are indicated in pink (S212, S216, S220) and in multisite motif-2 are indicated in red (T231, S235, S239, S243). **B.** The deletion mutant lacking both motif-1 and motif-2 (eIF6-CΔ36) was generated by substitution of T210 with a stop codon. Myc-eIF6-WT or myc-eIF6-CΔ36 were transfected into HCT116 cells, and 24 hours later, cells were briefly serum starved in 0.1% FBS for 4 hours. For the *in vitro* kinase assay, myc-eIF6-WT or myc-eIF6-CΔ36 was immunoprecipitated and incubated in the presence or absence of recombinant GSK3. *In vitro* kinase reactions were then subjected to autoradiography and western blotting using anti-myc antibody. Each experiment was repeated three independent times. Asterisks indicate the presence of a doublet as seen by autoradiography and western blotting. **C.** Mass Spectral (MS) analysis was carried out on myc-eIF6 immunoprecipitated from serum starved HCT116 cells that were incubated with or without recombinant GSK3. Samples were excised from coomassie-stained acrylamide gels and analyzed by *nano*LC-MS/MS analysis. Plot shows the MS/MS spectrum along with matched fragmentation mass table showing specific T231 phosphorylation in the region of interest.

We then determined if the immunoprecipitated eIF6 is directly phosphorylated by recombinant GSK3. Our results show that eIF6 is indeed phosphorylated by GSK3β *in vitro* and was specific since we did not observe a similar phosphorylated form in the empty vector control lanes (Fig. 1B, S1C). We also observed autophosphorylation of GSK3β, which has been reported before^44^.

We observed a doublet of immunoprecipitated myc-eIF6 in the western blot and this upward shift or slower migration is highly likely to represent a post-translationally modified form. The intensity of the doublet was found to be quite variable between experiments and was not cell-type specific. The doublet is unlikely to be a cleavage product since the myc-tag antibody that recognizes the N-terminus and the eIF6 antibody that is specific to the C-terminal tail were able to detect both bands. Analysis of the *in vitro* kinase assay revealed two phosphorylated forms of eIF6, which could indicate a basal and a hyperphosphorylated form (Fig 1B, S1C). We further tested for specificity of phosphorylation by eliminating the two multisite motifs in the C-terminus by creating a deletion mutant that lacks the last 36 amino acid residues (eIF6-ΔC36). GSK3β did not phosphorylate eIF6-ΔC36 whereas it phosphorylated the full-length eIF6 suggesting that the last 36 residues are critical for phosphorylation (Fig. 1B). We did observe a faint phosphorylated form in the eIF6-ΔC36 lane (Fig. 1B). Prolonged exposures of the blots have captured a similar non-specific band in the myc-empty vector lane incubated with the kinase (Fig. S1D). This residual background could be associated with immunoprecipitation or some non-specific phosphorylation observed in the absence of sites that are specific to the kinase.

### GSK3β phosphorylates multiple sites within the last 20 amino acid residues of eIF6

To identify the specific residues phosphorylated by GSK3β, we carried out nanoscale liquid chromatography with tandem mass spectrometry analysis (*nano*LC-MS/MS) on the immunoprecipitated eIF6 sample incubated with and without GSK3β. MS analysis revealed that the T231 residue was phosphorylated by GSK3β (Fig. 1C, S1E, F). The relative abundances of the peptides in the samples incubated with and without the kinase were found to be identical (Fig. S2A, B). However, we were unable to detect other sites of phosphorylation among the last 36 residues in the C-terminal tail. This could be attributed to either a better preservation of the T231 phosphorylation relative to other sites or the limitations of MS analysis to capture two or more sites of phosphorylation within a single peptide using immunoprecipitated sample. Phosphorylation of the T231 residue indicated that motif-2 in the C-terminal tail is phosphorylated by GSK3β. Motif-2 is in the disordered region of the C-terminal tail of eIF6 that is predicted to protrude outside the core structure, which would be favorable for regulatory interactions^10,16^. Also, all the predicted sites indicated in motif-2 were previously identified to be modified by phosphorylation in global proteomic studies including those performed under cellular conditions of stress^35–41^. Furthermore, previous studies did not detect phosphorylated residues in motif-1 of mammalian eIF6 (PhosphoSitePlus). To test if motif-2 is critical for recognition by GSK3β, a deletion mutant lacking the last 20 residues in the C-terminal tail (eIF6-ΔC20) was generated. Indeed, deletion of just motif-2 greatly reduced eIF6 phosphorylation, which in combination with the MS analysis indicated that the sites of phosphorylation are in motif-2 (Fig. 2A).

**Figure 2.**
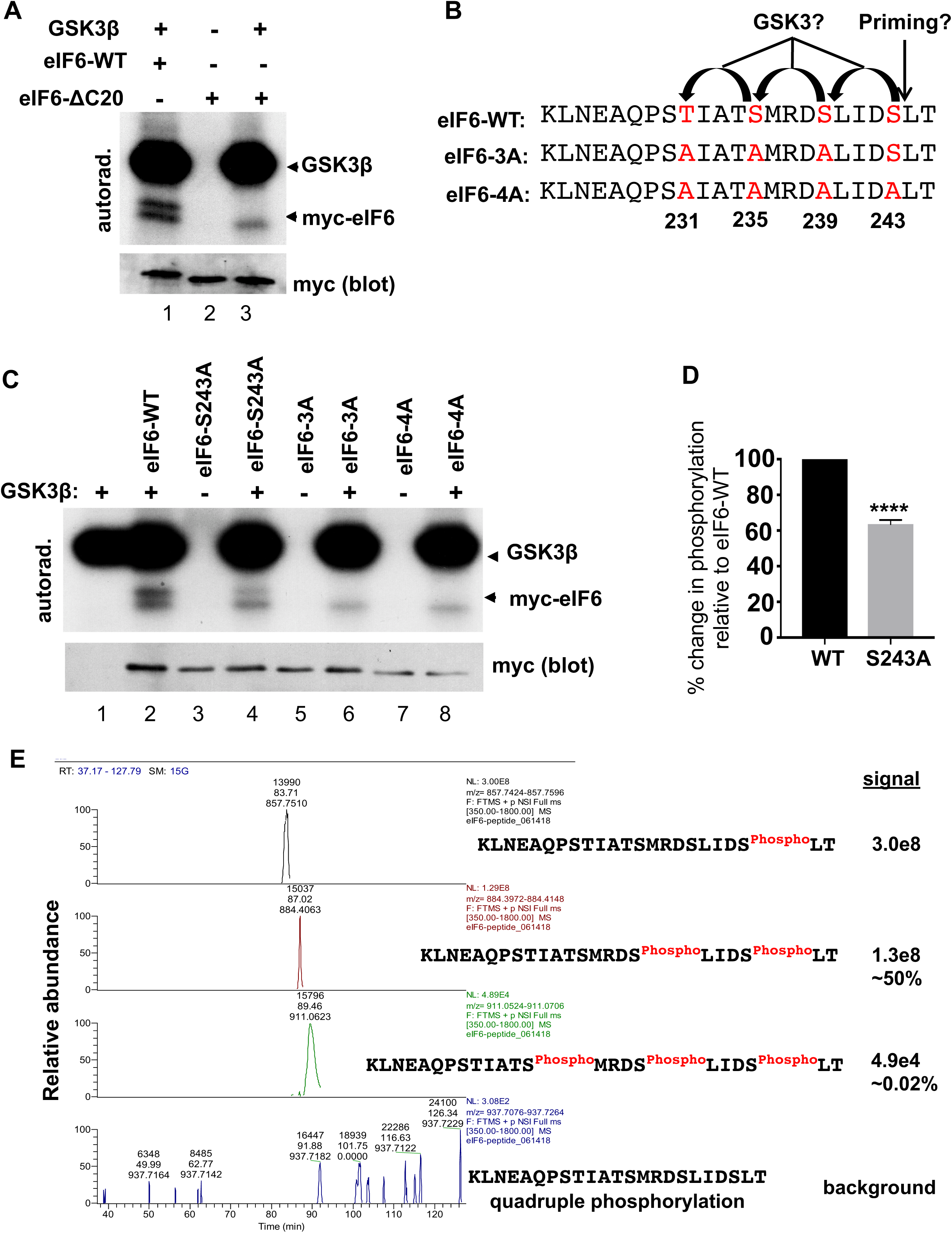
GSK3β phosphorylates multiple sites of eIF6 in a sequence. **A.** The deletion mutant lacking just motif-2 (eIF6-CΔ20) was generated by substitution of E226 with a stop codon. Myc-eIF6-WT or myc-eIF6-CΔ20 was immunoprecipitated from serum starved HCT116 cells, and the *in vitro* kinase reactions were assayed by autoradiography and western blotting. Each experiment was repeated three independent times. **B.** For the eIF6-3A and eIF6-4A mutants, the specific phospho-sites that were substituted with alanines are indicated and compared to the sequence of eIF6-WT. **C.** Myc-eIF6-WT or myc-tagged phospho-site mutants were immunoprecipitated from HCT116 cells that were serum starved for 4 hours. *In vitro* kinase reactions were carried out in the presence and absence of recombinant GSK3β and were analyzed by autoradiography and by western blotting using anti-myc antibody. Each experiment was repeated three independent times. **D.** Graph represents the percent change in the phosphorylated levels of the eIF6-S243A phospho-site mutant relative to the eIF6-WT as determined from the autoradiographs indicated in Fig. 2C. Values represent the SEM of three independent experiments. Asterisks indicate significantly different as determined by an unpaired two-tailed t-Test: p<0.0001. **E.** For *nano*LC-MS/MS analysis, an eIF6 phosphopeptide (23 amino acids) carrying a phosphate at S243, the putative priming site, was synthesized (Aapptec) and incubated with recombinant GSK3β. Extracted ion chromatograms show relative abundance levels of singly (pre-charged) and multiple residues phosphorylated by GSK3β in the region of interest.

To further identify the specific sites of phosphorylation in motif-2, we generated phospho-site mutants where individual serine or threonine residues within motif-2 were substituted with alanine (Fig. 2B). Interestingly, substitution of the potential priming site serine-243 with alanine resulted in a significant 40% reduction in phosphorylation in comparison to the wild type eIF6 (eIF6-WT) (Fig. 2C, D). This suggested that in the absence of the S243 priming site, the S239, S235 and T231 sites are phosphorylated, however priming phosphorylation at S243 is required for efficient phosphorylation of eIF6. To determine if the other three sites are phosphorylated in a GSK3-specific manner, we generated a 3A phospho-site version carrying the following serine to alanine substitutions: T231A, S235A, S239A (Fig. 2B). In the absence of the T231, S235 and S239 sites, phosphorylation of eIF6 was greatly reduced with a complete loss of the hyperphosphorylated form suggesting that these are the key sites of phosphorylation (Fig. 2C). The loss of the eIF6 doublet that we normally see with the eIF6-WT protein in the autoradiograph (Fig. 1C, 2A) suggests that the slower migrating species is highly likely a hyperphosphorylated version. This shift in migration was consistent with the observation that the phosphomimic eIF6-4E mutant, where all 4 sites of phosphorylation were substituted with glutamate, migrated 1-2kDa higher on the gel (Fig. S3A). Similar results were obtained with the 4A mutant wherein all 4 sites were substituted with alanine, suggesting that all the four residues are key for phosphorylation by GSK3β and that S243 is the priming site (Fig. 2C).

### GSK3β phosphorylates multiple sites on eIF6 in a sequence

In the presence of multiple and adjacent recognition sites, GSK3 phosphorylates residues in a sequence starting from the C-terminus. Examples of sequential phosphorylation include substrates such as β-catenin and glycogen synthase that carry 3 and 4 GSK3-specific sites respectively^57–61^. To determine if phosphorylation of eIF6 is also sequential, we carried out MS analysis on a phosphopeptide incubated with GSK3β *in vitro*. The phosphopeptide was designed to carry a phosphate group on the S243 site (priming site) to mimic a primed state (Fig. 2E). MS analysis revealed that GSK3β phosphorylated a major fraction of the peptides at the S239 site, the first site in the sequence. Also, peptides phosphorylated at both the S235 and S239 sites (Fig. 2E, S3B) were detected. Detection of quadruple phosphorylation was at or below baseline (Fig. 2E). However, the T231 site was identified earlier using our previous MS analysis on immunoprecipitated eIF6 (Fig. 1C). Interestingly, we were unable to detect any doubly phosphorylated-S235/S243 or T231/S243 peptides, which further suggested that phosphorylation occurs in a sequence and has to include the S239 site. These MS results in combination with eIF6-3A and eIF6-4A mutants strongly suggested that the phosphorylation of eIF6 by GSK3β occurs in a sequence starting with the S239 site, followed by the S235 site and ending with the T231 site.

### Priming is critical for robust phosphorylation of eIF6

To ascertain the significance of priming, MS analysis was carried out on an unprimed C-terminal peptide (non-phosphorylated) incubated with GSK3β *in vitro* (Fig. S4). In the absence of priming, phosphorylation could be barely detected above baseline for the S243 site and S239 sites (Fig. S4). Only 0.2% of the total were phosphorylated at the S243 site, and 0.002% of the total were phosphorylated at the S239 site (Fig. S4). These results strongly indicated that priming is critical for robust phosphorylation of eIF6.

We further performed the kinase assay using a priming-recognition site mutant of GSK3β (R96A) that is unable to phosphorylate substrates that require priming^45,54,55^. The arginine-96 to alanine substitution in GSK3β disrupts the priming recognition site within the kinase^45,54^. Substrates that are primed by phosphorylation, dock on to this priming recognition site followed by phosphorylation at the catalytic site^45,54^. However, substrates that do not require priming are phosphorylated by the GSK3β-R96A mutant with efficiency similar to the wild type GSK3β^45,54,55^. In our experiments, as expected, wild type GST-GSK3β phosphorylated eIF6 (Fig. 3A). However, no such doublet was observed with the GST-GSK3-R96A mutant suggesting that recognition of the primed phosphorylation is critical for efficient phosphorylation of eIF6 (Fig. 3A).

**Figure 3.**
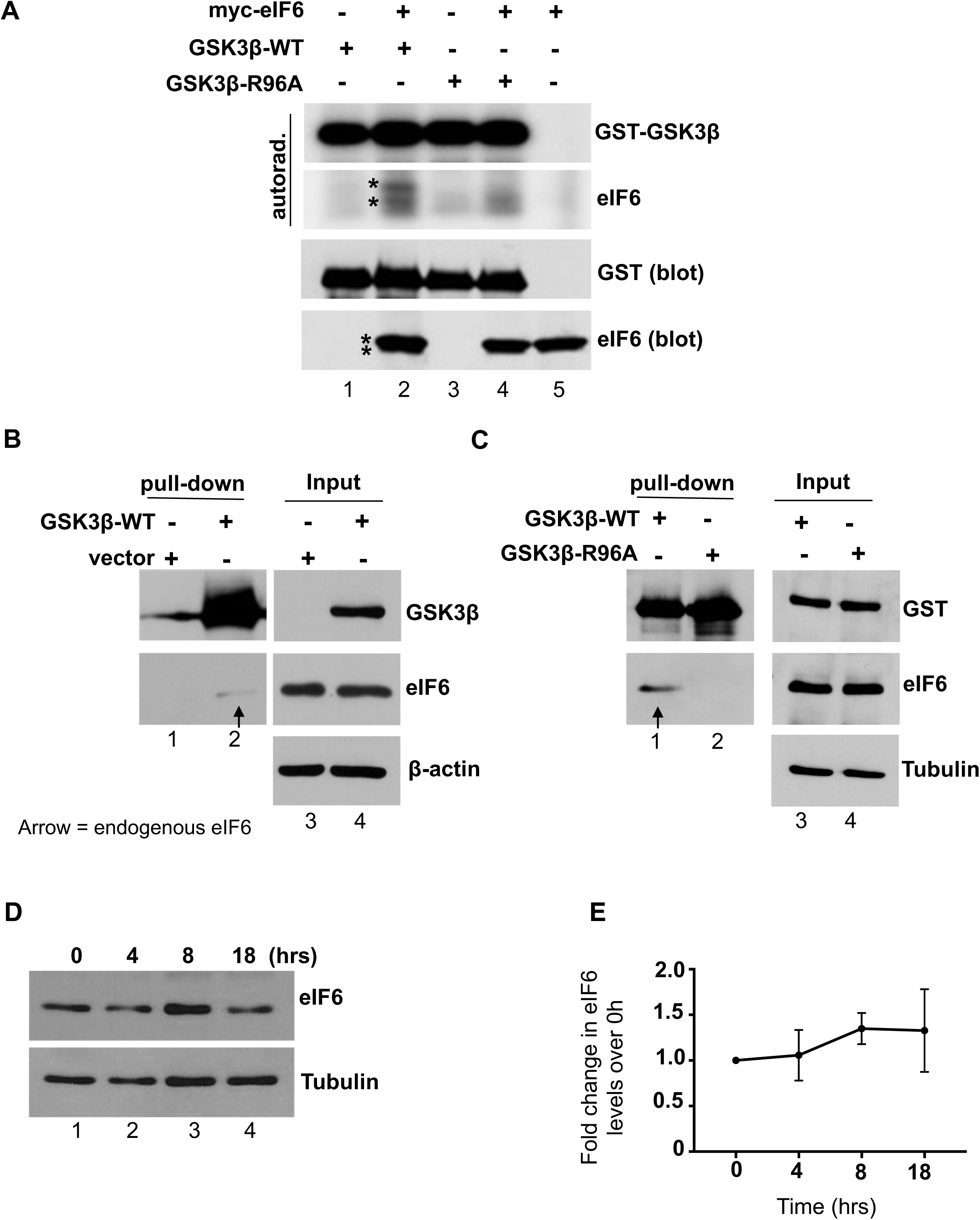
Priming dependent phosphorylation of eIF6 by GSK3β. **A.** GST-GSK3β-WT, GST-GKS3β-R96A mutant, or myc-eIF6 were transfected into 293T cells and 24 hours later, cells were briefly serum starved for 4 hours. GST-GSK3β-WT or GST-GKS3β-R96A was pulled-down and eluted with glutathione as described in methods. Myc-eIF6 was immunoprecipitated from 293T cells. GST-GSK3β-WT or GST-GKS3β-R96A was incubated in the presence and absence of myc-eIF6 for the *in vitro* kinase reactions. Kinase reactions were screened by autoradiography and by western blotting. Blots were probed with anti-eIF6 and anti-GST antibodies. Each experiment was repeated three independent times. Asterisks indicate the presence of a doublet as seen by autoradiography and western blotting. **B and C.** To capture interactions between GST-GKS3β and endogenous eIF6, HCT116 cells were transfected with GST-GKS3β-WT or empty vector (**B**), or with GST-GKS3β-WT or GST-GKS3β-R96A mutant (**C**). 24 hours after transfection, cells were serum starved for 4 hours. Blots were probed with anti-GKS3β, anti-eIF6 and anti-GST antibodies. Each experiment in B and C was repeated three and two independent times respectively. Whole-cell lysates (inputs) of samples used for the pull-downs were analyzed by western blotting and probed with the indicated antibodies (**lanes 3, 4**). β-Actin and αTubulin were used as the loading control. **D.** Lysates of HCT116 cells were collected just prior to serum starvation at 0 hours and again at 4, 8, and 18 hour time points. Samples were analyzed by western blotting and blots were probed with anti-eIF6 antibody and anti-αTubulin antibody (loading control) (n=3). **E.** Total eIF6 protein levels were quantitated using the blots represented in **D**. Values represent the standard error of the mean (SEM) of three independent experiments. All time points were compared with the 0 hour time point and no significant differences were found as determined by one-way ANOVA (Dunnett’s multiple comparison test).

### Phosphorylation of eIF6 in response to serum starvation in vivo

We also determined if eIF6 is phosphorylated in response to serum starvation *in vivo* by mass spectrometry. *nano*LC-MS/MS analysis revealed that the C-terminal tail of eIF6 carries a phosphorylation site that matched to both the S243 site (Fig. S5A) and S239 site (Fig. S5B). This could be attributed to a mix of peptides that were phosphorylated either at the S243 site or the S239 site. This indicates that the C-terminal tail of eIF6 is phosphorylated *in vivo* under serum-starved conditions.

### GSK3 interacts with endogenous eIF6 in human cells

Since GSK3 is activated under serum-starved conditions, we next tested if GSK3β interacts with eIF6 in human cells. GST-GSK3β pulled-down from HCT116 cells was found to interact with endogenous eIF6 (Fig. 3B, C). The interaction was found to be specific to wild type-GST-GSK3β as no such interaction was observed with either empty vector or with the priming-site recognition mutant (GST-GSK3β-R96A) (Fig. 3B, C). These results clearly indicate that GSK3β does interact with eIF6 *in vivo*.

### Altered sub-cellular localization of eIF6 in response to starvation is regulated by GSK3

We then wanted to determine the effect of GSK3-dependent phosphorylation on eIF6 levels or localization. We observed that the total levels of eIF6 were not altered by overexpression of GST-GSK3β, suggesting that GSK3β may not regulate the total protein levels. This was also confirmed by probing for eIF6 protein levels in response to long-term serum starvation when GSK3β is fully active (Fig. 3D, E). We did not detect significant changes in total eIF6 protein levels even after 18 hours of starvation suggesting that GSK3β does not regulate the stability or synthesis of eIF6 protein (Fig. 3D, E).

Since the total levels of eIF6 were unaltered, we next tested whether GSK3β regulates eIF6 function by altering its subcellular localization. We also tested if long-term serum starvation that correlates with complete activation of GSK3β altered the subcellular localization of eIF6.

Immunofluorescence staining of endogenous eIF6 showed that eIF6 localization to the cytoplasm was enhanced in response to serum starvation in HCT116 cells (Fig. 4A). This was also observed in HeLa cells where serum starvation caused cytoplasmic (punctate) accumulation of eIF6, which was reversed within 2 hours by the re-addition of serum indicating that the response is prompt (Fig. 4B). These results were further confirmed by using a different monoclonal antibody specific for eIF6, which ruled out non-specific staining associated with the antibody (Fig. S6A). To further determine if this altered sub-cellular localization was dependent on GSK3 activity, cells were treated with CHIR99021 and SB415286, two highly selective GSK3-specific inhibitors^62,63^. The potency of the GSK3-specific inhibitors was validated by probing for a loss of phosphorylation of β-catenin at GSK3-specific sites (S33/S37/T41)^57–59,64^. Inhibitor treatment resulted in greater than 50% loss in β-catenin phosphorylation suggesting that GSK3 is inhibited under these conditions (Fig. S6B, C). Treatment with the GSK3-specific inhibitors, eIF6 localization to the nucleus was rescued (Fig. 4A). Similar results were also found in a normal rat intestinal epithelial cell line (RIE-1) where eIF6 shows cytoplasmic accumulation in response to serum starvation, which is reversed by inhibition of GSK3 (Fig. S6D).

**Figure 4.**
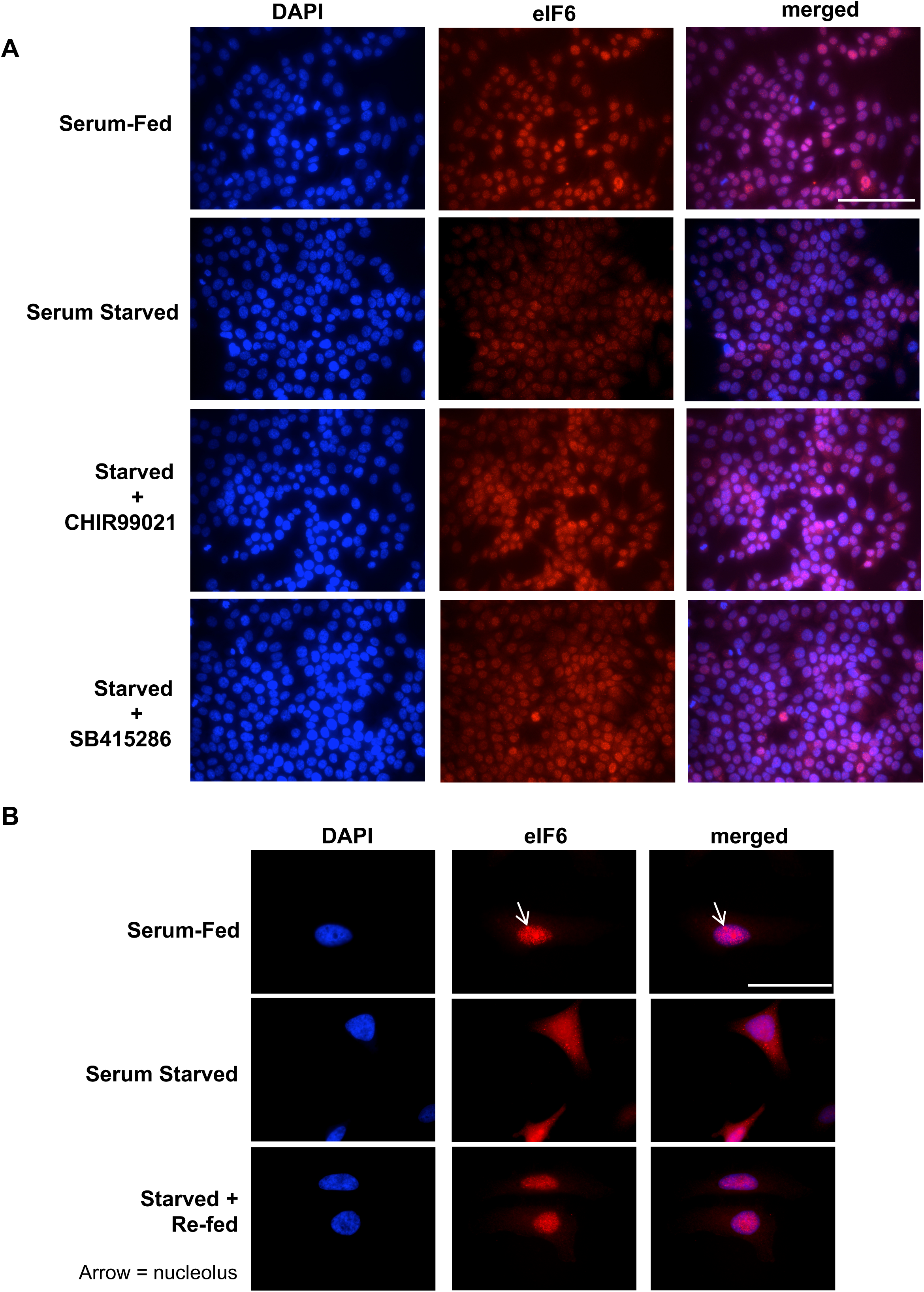
Altered subcellular localization of eIF6 in response to serum starvation is regulated by GSK3. **A.** HCT116 cells were serum starved for 24 hours, and were treated for with either vehicle (0.1% DMSO), 10μM CHIR99021or 25μM SB415286 for 3 hours. Cells fed with 10% FBS and treated with vehicle were used as the non-starved (serum-fed) controls. Cells were fixed and stained for endogenous eIF6 (red) using anti-eIF6 monoclonal antibody (Cell Signaling Technology) and nuclei were stained with DAPI (blue) and analyzed by immunofluorescence microscopy. Each experiment was repeated three independent times. Scale bar =100μm. **B.** HeLa cells were serum-fed or serum starved for 24 hours as indicated in **A**. For re-feeding, serum starved cells were fed with 10% FBS for 2 hours. Cells were fixed and stained for endogenous eIF6 (red) using anti-eIF6 monoclonal antibody (Cell Signaling Technology) and nuclei were stained with DAPI (blue) and analyzed by immunofluorescence microscopy. Each experiment was repeated three independent times. Scale bar = 50μm.

We further quantitated eIF6 levels in the nuclear and cytoplasmic fractions by subcellular fractionation. The purity of different fractions was confirmed by probing for Topoisomerase IIβ, a nuclear marker and αTubulin, a cytoplasmic marker. eIF6 was distributed almost equally between the nuclear and cytoplasmic fractions in serum-fed controls (53% nucleus, 47% cytoplasm), and no significant difference was found between the two fractions (Fig. 5A, B). Similar to the imaging results, we found that in the serum starved cells there was a significant decrease in eIF6 levels in the nucleus with a corresponding increase in the cytoplasm (∼32-33% in nucleus, ∼67-68% in cytoplasm) (Fig. 5A, C, D, E). Inhibition of GSK3 completely restored the nuclear-cytoplasmic distribution as seen in the CHIR990021-treated cells (54% nucleus, 46% cytoplasm) (Fig. 5D, F).

**Figure 5.**
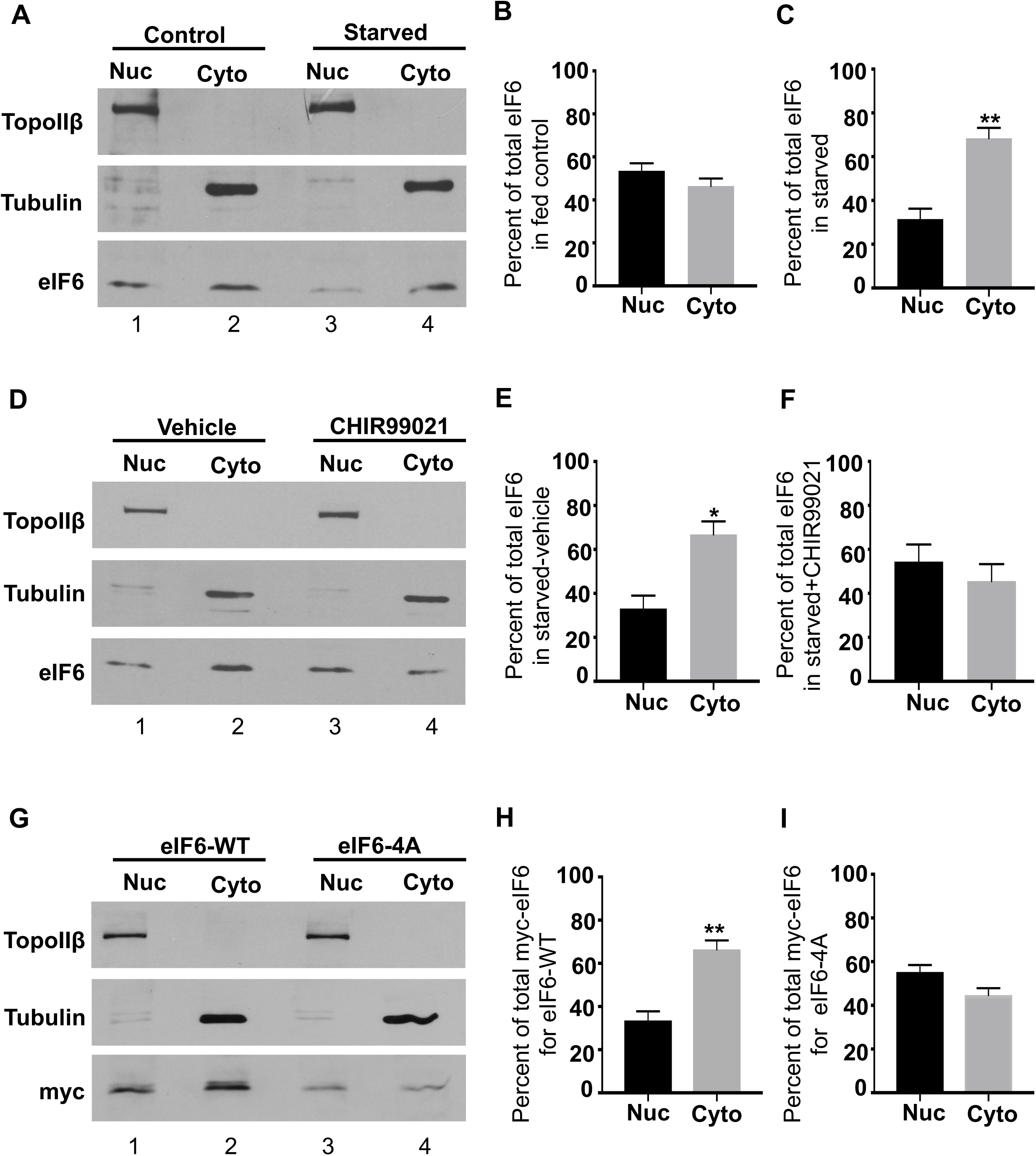
The phosphodead mutant does not exhibit enhanced cytoplasmic accumulation in response to serum starvation. **A, B and C.** For nuclear and cytoplasmic extraction, HCT116 cells were serum-fed (lanes 1, 2) or serum starved for 24 hours (lanes 3, 4) followed by subcellular fractionation. Samples were analyzed by western blotting. Blots were probed with anti-eIF6, anti-αTubulin (cytoplasmic marker), and anti-Topoisomerase IIβ antibodies (nuclear marker) (**A**). Blots represented in **A**., were quantitated and values represent the SEM of four independent experiments (**B and C**). Percent of nuclear and cytoplasmic fractions relative to the total (sum of nuclear and cytoplasmic eIF6 levels) are plotted. Difference in eIF6 levels between the nuclear and cytoplasmic fractions for non-starved controls (**B**) were not significant as determined by an unpaired two-tailed t-Test but were found to be significant for the starved sample (**C**) and asterisks indicate: p=0.0015. **D, E and F.** HCT116 cells were serum starved for 24 hours and treated with vehicle (lanes 1, 2) or 10μM CHIR99021 (lanes 3, 4) (5 hours) followed by subcellular fractionation. Samples were analyzed by western blotting and blots and probed with the indicated antibodies (**D**). Blots represented in **D**., were quantitated and values represent the SEM of three independent experiments (**E and F**). Percent of nuclear and cytoplasmic fractions relative to the total (sum of nuclear and cytoplasmic eIF6 levels) are plotted. The difference in eIF6 levels between the nuclear and cytoplasmic fractions for the starved-vehicle control (**E**) was found to be significant as determined by an unpaired two-tailed t-Test and asterisk indicates: p=0.015, but no significant difference was found between the CHIR99021-treated fractions (**F**). **G, H and I.** Myc-tagged eIF6-WT or eIF6-4A mutant were transiently expressed in HCT116 cells stably knocked-down for eIF6 (eIF6-KD) and cells were serum starved for 24 hours followed by subcellular fractionation. Samples were analyzed by western blotting and blots were probed with anti-myc, anti-αTubulin, and anti-Topoisomerase IIβ antibodies (**G**). Blots represented in **G**., were quantitated and values represent the SEM of three independent experiments (**H and I**). Percent of nuclear and cytoplasmic fractions relative to the total (sum of nuclear and cytoplasmic eIF6 levels) are plotted. The difference in myc-eIF6 levels between the nuclear and cytoplasmic fractions for eIF6-WT (**H**) was found to be significant as determined by an unpaired two-tailed t-Test and asterisks indicates: p=0.0052, but no significant difference was found for the eIF6-4A mutant (**I**).

To determine the significance of phosphorylation in regulating the subcellular localization of eIF6 in response to starvation, we generated myc-tagged phosphodead mutant of eIF6 (eIF6-4A) where all 4 sites of phosphorylation were substituted with alanine. We also generated the myc-tagged eIF6-ΔC mutant that lacked the last 20 amino acid residues in the C-terminus including the GSK3-specific sites of phosphorylation. To analyze the effect of myc-tagged wild type and mutants in the absence of endogenous eIF6, they were expressed in a cell line that was stably knocked down for eIF6 (eIF6-KD) (Fig. S7A). Endogenous eIF6 levels were reduced to greater than 85% by using shRNA targeted against eIF6 (Fig. S7B). Analysis of the subcellular localization of the phosphodead mutant revealed that eIF6-4A does not accumulate in the cytoplasm in response to starvation unlike the WT cells (Fig. 5G, H, I). To determine if eIF6-ΔC localization was similar to the eIF6-4A mutant, we analyzed cells that stably express either the eIF6-ΔC mutant or eIF6-WT (Fig. S7C). The eIF6-ΔC mutant did not exhibit significant cytoplasmic accumulation in response to starvation unlike the WT (Fig. S8A, B, C). The total input levels of eIF6 under all conditions tested are shown in Fig. S8D, E, and F. The nuclear cytoplasmic distribution of the eIF6-4A mutant was not found to be different under fed conditions (Fig. S8G). These results strongly suggest that eIF6 exhibits cytoplasmic accumulation in response to serum starvation and this altered localization is regulated by GSK3-dependent phosphorylation of the C-terminal tail.

### Physiological significance of phosphorylation in response to serum starvation

We performed polysome profile assay to determine the physiological effect of disrupting the GSK3-specific sites of phosphorylation. Polysome profiles of starved (wild type control) cells in comparison to the serum-fed cells showed that serum starvation causes a profound inhibition of translation as shown by an increase in the levels of free 60S, 40S and 80S subunits with a concomitant decrease in the heavier polysome levels (Fig. S9A) and this was consistent with previous reports. Interestingly, under serum starved conditions, the eIF6-4A mutant exhibited a significant 20% to 25% increase in the levels of free 60S subunits (Fig. 6A, B). Enhanced 60S levels were also observed for the eIF6-ΔC mutant under serum starved conditions (Fig. 6C). A consistent increase in the levels of 80S monosomes along with a modest and consistent decrease in heavier polysome levels under starved conditions was observed for both the eIF6-4A and the eIF6-ΔC mutant cells (Fig. 6A, B, C). These results suggest that translation is further inhibited in the mutant cells under starved conditions.

**Figure 6.**
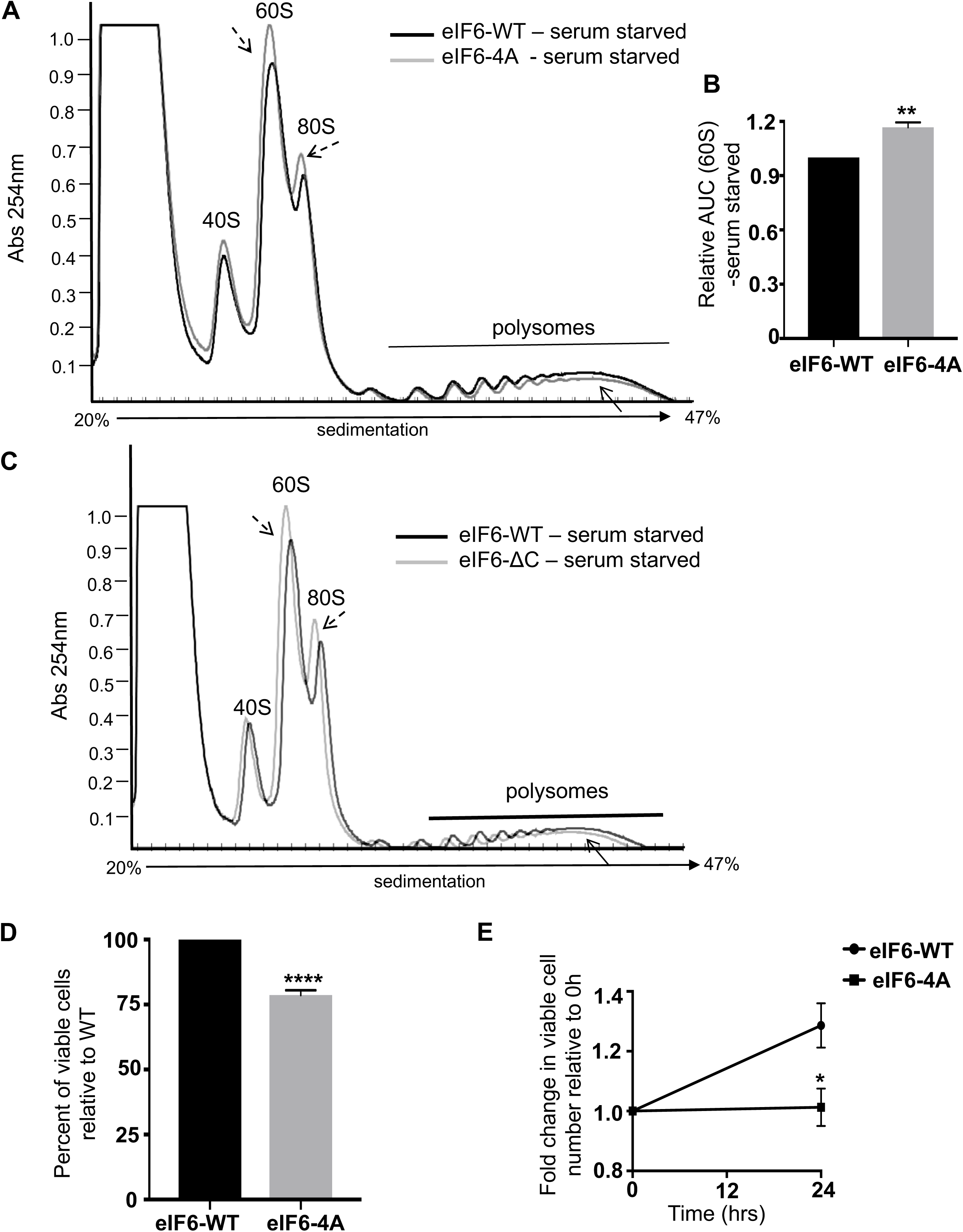
Polysome profile analysis of the phosphodead mutant of eIF6 shows enhanced inhibition of translation in response to serum starvation. **A and B.** Representative polysome profile of serum starved HCT116 cells stably expressing myc-tagged eIF6-4A mutant was overlaid with the eIF6-WT profile. Cells were serum starved for 24 hours prior to analysis. (**A**). Area Under a Curve (AUC) of 60S peak for eIF6-4A mutant normalized to AUC of the 60S peak of eIF6-WT is indicated in the bar graph (**B**). Graph represents the standard error of the mean (SEM) of three independent experiments. Asterisks indicate significant differences between eIF6-WT and eIF6-4A with p=0.0035 as determined by an unpaired two-tailed t-Test. **C.** Representative polysome profile of HCT116 cells stably expressing myc-tagged eIF6-ΔC mutant was overlaid with the eIF6-WT profile. Cells were serum starved for 24 hours prior to analysis. Profiles are representative of three independent experiments. **D.** Bar graph represents the percentage of viable HCT116 cells stably expressing eIF6-4A mutant relative to eIF6-WT. Cells were serum starved for 24 hours and subjected to trypan blue dye exclusion assay. Graph represents the standard error of the mean (SEM) of three independent experiments. Asterisks indicate significant differences between eIF6-4A and eIF6-WT with p<0.0001 as determined by an unpaired two-tailed t-Test. **E.** Plot represents the fold change in viable HCT116 cells stably expressing eIF6-4A mutant relative to eIF6-WT. Cells were serum starved for 24 hours and subjected to MTS assay. Plot represents the standard error of the mean (SEM) of three independent experiments. Asterisk indicates significant differences between eIF6-4A and eIF6-WT with p=0.0131 as determined by an unpaired two-tailed t-Test.

We then determined if altered translation profile in the eIF6-4A mutant affected cell growth or survival in response to starvation. As shown in Fig. 6D and 6E, there was a significant 20% to 25% decrease in the number of viable eIF6-4A mutant cells in comparison to eIF6-WT. No difference in viable cell number was observed between eIF6-WT versus eIF6-4A mutant under serum-fed conditions (Fig S9B). Analysis of the MTS assay further indicated that even in the absence of serum, cell proliferation was not fully inhibited for eIF6-WT but was completely stalled for the eIF6-4A mutant (Fig. 6E). These results indicate that the eIF6-4A mutant significantly impacts cell growth in response to starvation.

## Discussion

The metabolic response to starvation is to conserve energy and limit processes with high-energy requirements such as translation and ribosome biogenesis^22–31^. Here, we provide a novel link between starvation response and regulation of both translation initiation and ribosome availability through the control of translation initiation factor eIF6. Despite its extensive role in metabolism, eIF6 function in starvation-response is poorly understood. We show for the first time that eIF6 is regulated in response to starvation-induced stress and that the regulation hinges on the GSK3 signaling pathway that is prolifically active under such nutrient-deprived conditions. This study also identifies GSK3 as one of the kinases that phosphorylates the multiple sites in the C-terminal tail of eIF6. Although several studies have shown that the C-terminal tail of eIF6 is heavily phosphorylated, the identity of the kinases involved, other than PKCβII, were largely unknown^12^. Our results show that GSK3 sequentially phosphorylates the C-terminal tail of eIF6 on three sites (S239, S235, and T231) and efficient phosphorylation of these sites requires priming at S243. It is currently unclear as to which priming kinase works in concert with GSK3 and the identity and functional significance of this priming kinase will be investigated in future studies.

We and others have shown that in response to nutrient deprivation, protein synthesis is inhibited with a decrease in polysome levels along with a corresponding increase in free 60S subunits and inactive 80S monosomes^22–33^. Any changes to the 60S or 40S levels can drastically impact cell growth as shown for several ribosomopathies. Our results indicate that the phosphorylation of the C-tail of eIF6 and its cytoplasmic accumulation is important to maintain a basal level of protein synthesis under starved conditions by balancing the free pool of 60S and inactive 80S monosomes.

Interestingly, one of the sites of phosphorylation, the S235 site, is also phosphorylated by the RACK1-PKCβII complex^12^. Biochemical studies show that the RACK1-PKCβII-dependent phosphorylation of eIF6 results in its release from the 60S subunits and enables active 80S formation^12^. Substitution of the serine-235 site with alanine blocks eIF6 release and inhibits active 80S formation^12^. Apart from *in vitro* studies, the S235 site has also been shown to be important for normal growth and translation *in vivo*^17,18^. Studies in *eIF6*+/- MEFs reconstituted with the S235A mutant show that the S235 site is critical for eIF6 function in stimulating translation in response to insulin and phorbol esters^17–19^. It is not uncommon for a single site to be regulated by multiple kinases in a context-dependent manner. Since PKCβII is active under conditions of growth and proliferation, whereas GSK3 is active under growth-deprived conditions, the two kinases could phosphorylate the S235 site based on the polarizing cellular cues of growth and starvation. Regardless of the identity of the kinase, phosphorylation at the S235 site is expected to exert a similar effect and inhibit eIF6 association with the ribosome. However, in the case of eIF6 phosphorylation by GSK3β, additional sites are also phosphorylated, which could further perturb the local protein structure.

We observed enhanced cytoplasmic accumulation of eIF6 in response to serum starvation and inhibiting the GSK3-dependent phosphorylation of eIF6 rescues the subcellular distribution. Since GSK3 is predominantly cytoplasmic with a small nuclear fraction^40^, it is likely that phosphorylation by GSK3 leads to increased cytoplasmic retention of eIF6. A previous report in COS-7 cells indicates that the percentage of eIF6 in the cytoplasm is ∼70% compared to the ∼50% distribution that we observe in HCT116 cells^65^. In our study, the increased cytoplasmic accumulation of eIF6 in response to starvation was consistently observed among different cell lines including the normal RIE-1 cells and HeLa cells. A previous study showed that the nuclear import of eIF6 was mediated by calcium-activated calcineurin phosphatase whereas the nuclear export was mediated by phosphorylation of eIF6 at S174/175 residues by nuclear casein kinase1 in COS-7 cells^65^. Thus, our results and previous data suggest that altering the subcellular localization of eIF6 may be a predominant mode of regulating eIF6 function, and the underlying signaling mechanisms may vary based on the growth stimulus and stressed states.

Based on these results, we propose a model of eIF6 regulation in response to starvation (Fig. 7). Under nutrient or growth-limiting conditions, AKT and mTORC1-p70S6K signaling mechanisms are inactivated, which in turn activates GSK3 (Fig. 7). Activation of GSK3 in concert with a priming kinase leads to phosphorylation of eIF6 at four sites. Phosphorylation by GSK3 enhances the cytoplasmic accumulation of eIF6, which is critical to maintain the free pool of 60S subunits and aids in cellular adaptation to starvation. The free pool of 60S could be maintained by altering 60S biogenesis or maturation. These results indicate that eIF6 contributes to the starvation-induced attenuation of global protein synthesis in concert with other adaptive mechanisms such as inhibition of mTOR, and GSK3-dependent inhibition of eIF2B.

**Figure 7.**
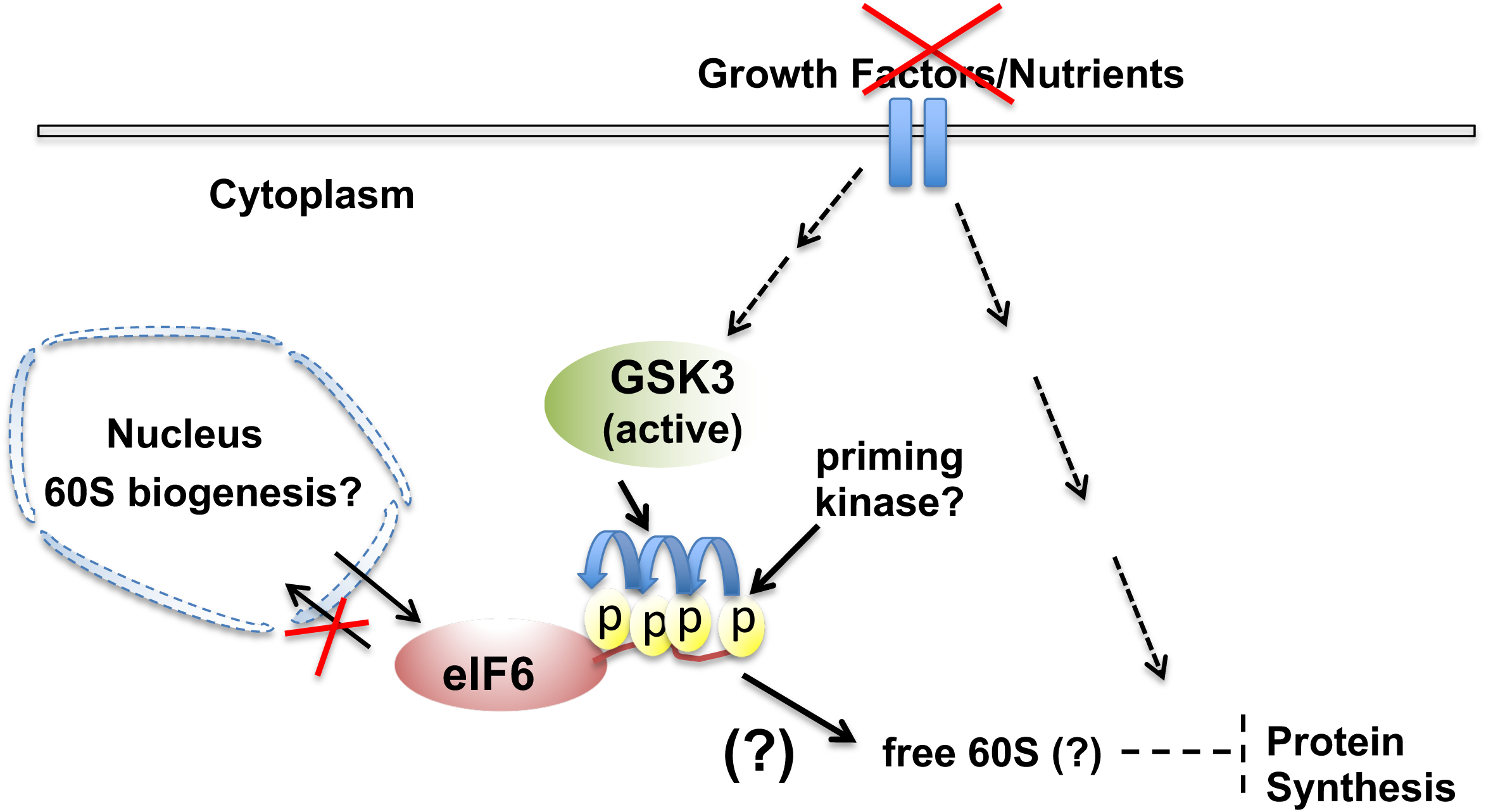
Proposed model for regulation of eIF6 by GSK3. In response to serum starvation or in the absence of nutrients or growth factor stimulation, GSK3 is potently activated by the inhibition of AKT and mTOR signaling pathways. Active GSK3 phosphorylates eIF6 on three specific sites, which is potentiated by priming at the C-terminal site by an unidentified kinase. eIF6 exhibits cytoplasmic accumulation in response to stress induced by serum starvation, which is regulated by GSK3. Phosphorylation regulates 60S levels and translational response to starvation, which affects cell growth. It is currently unclear as to how phosphorylation of eIF6 maintains free 60S subunit pools (maturation/biogenesis?) and how it contributes to the starvation-induced attenuation of global protein synthesis in concert with other adaptive mechanisms such as inhibition of mTOR, and GSK3-dependent inhibition of eIF2B.

## Acknowledgements

We thank Dr. Grzegorz Sabat and Dr. Greg Barrett-Wilt at the University of Wisconsin-Madison Mass Spectrometry Core for helping us with generation and analysis the mass spectrometry data. We also thank Dr. Edwin Antony for his intellectual input and for providing initial resources for the work. This study is supported by the NIH-R15GM119103 grant provided to CP. This study is supported by the NIH-GM126477 grant, by the Wisconsin Women’s Health Foundation through the Markos Family Breast Cancer Woman Faculty Scholar grant, and start-up funds provided by Marquette University to SO.

## Author contributions

The study was conceived by SO, who designed and performed some of the preliminary experiments and critically analyzed the results generated by CJ, JE and DM, and wrote the manuscript. CJ performed most of the experiments, interpreted the results and generated the associated images. CJ, JE, and DM wrote the methods section of the manuscript, which was edited by SO. CP generated the GSK3 constructs and contributed to the design of the priming site mutant experiments and critically reviewed the manuscript. DM performed the immunofluorescence experiments and analyzed the data. Majority of the clones were generated by JE, who designed some of the molecular cloning strategies and analyzed the data.

## Supplementary Figure Legend

**Figure S1.**
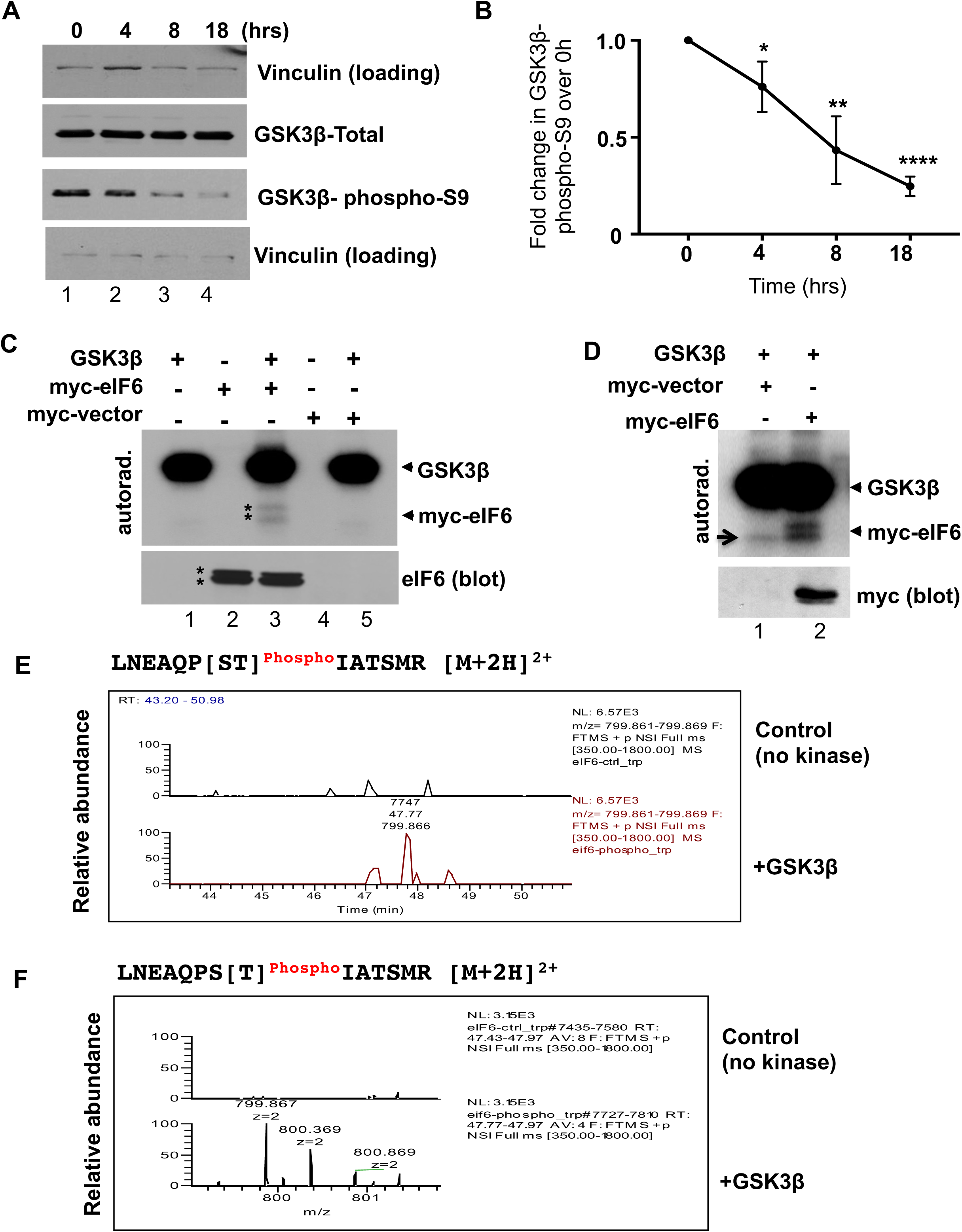
Activation of GSK3 in response to serum starvation. **A.** HCT116 cells were serum starved for 18 hours and cell lysates were collected just prior to serum starvation at 0 hours and at 4, 8 and 18 hours. Samples were analyzed by western blotting and blots were probed with anti-GSK3 (total), anti-GSK3-pS9 or anti-Vinculin antibody (loading control) (n=3). **B.** GSK3-Serine-9 phosphorylation levels were quantitated using the blots represented in A. Values represent the SEM of three independent experiments. Asterisks indicate significantly different as determined by an unpaired two-tailed t-Test. The difference in eIF6 levels between 0 hour versus 4 hours was significant with p=0.033, 0 hour versus 8 hours was significant with p=0.0008, and 0 hour versus 18 hours was significant with p=0.0001. **C.** HCT116 cells were transfected with myc-eIF6 or myc-empty vector, and 24 hours later, cells were briefly serum starved in 0.1% FBS for 4 hours. For the *in vitro* kinase assay, myc-eIF6 or myc-vector controls were immunoprecipitated and incubated in the presence or absence of recombinant GSK3. *In vitro* kinase reactions were then subjected to autoradiography and western blotting using anti-eIF6 antibody. Each experiment was repeated three independent times. Asterisks indicate the presence of a doublet as seen by autoradiography and western blotting. **D.** HCT116 cells were transfected with myc-eIF6-WT or myc-empty vector and subjected to immunoprecipitation followed by *in vitro* kinase reactions in the presence and absence of recombinant GSK3β as described previously (Fig. S1C). Arrow indicates the presence of a non-specific band observed for the myc-vector control (lane 1) after prolonged exposure (4 days). Each experiment was repeated three independent times. **E.** Identification of eIF6-T231 phosphorylation by MS/MS analysis. MS analysis of the peptide was carried out as described in Fig. 1D. Extracted Ion Chromatograms for the indicated unphosphorylated peptide signal is normalized to the highest. **F.** Mass spectrum of the indicated peptide shows normalized values averaged across extracted chromatogram.

**Figure S2.**
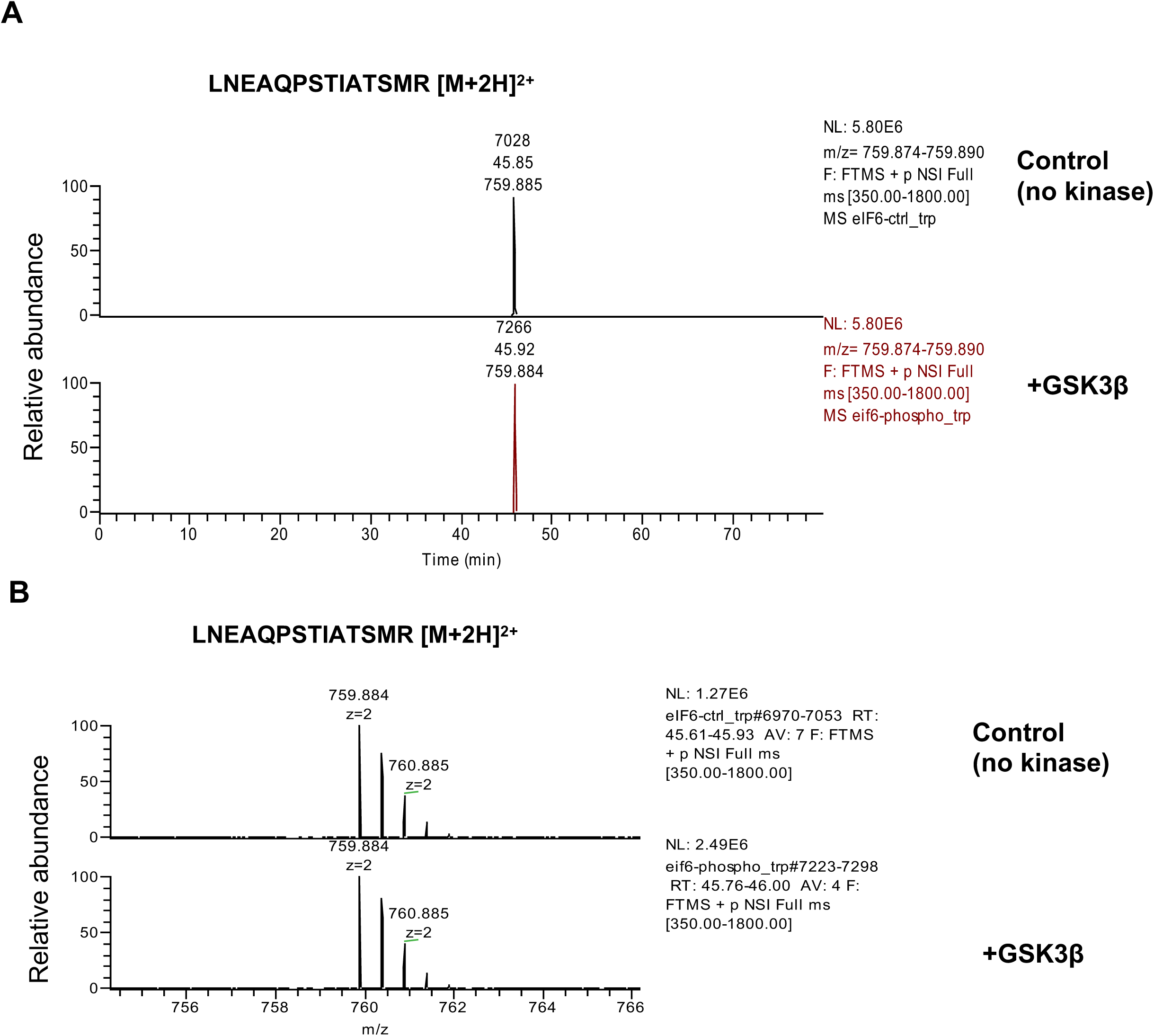
Identification of eIF6-T231 phosphorylation by MS/MS analysis. **A.** MS analysis of the peptide was carried out as described in Fig. 1D. Extracted Ion Chromatograms for the indicated unphosphorylated peptide signal is normalized to the highest. **B.** Mass spectrum of the indicated peptide shows normalized values averaged across extracted chromatogram.

**Figure S3.**
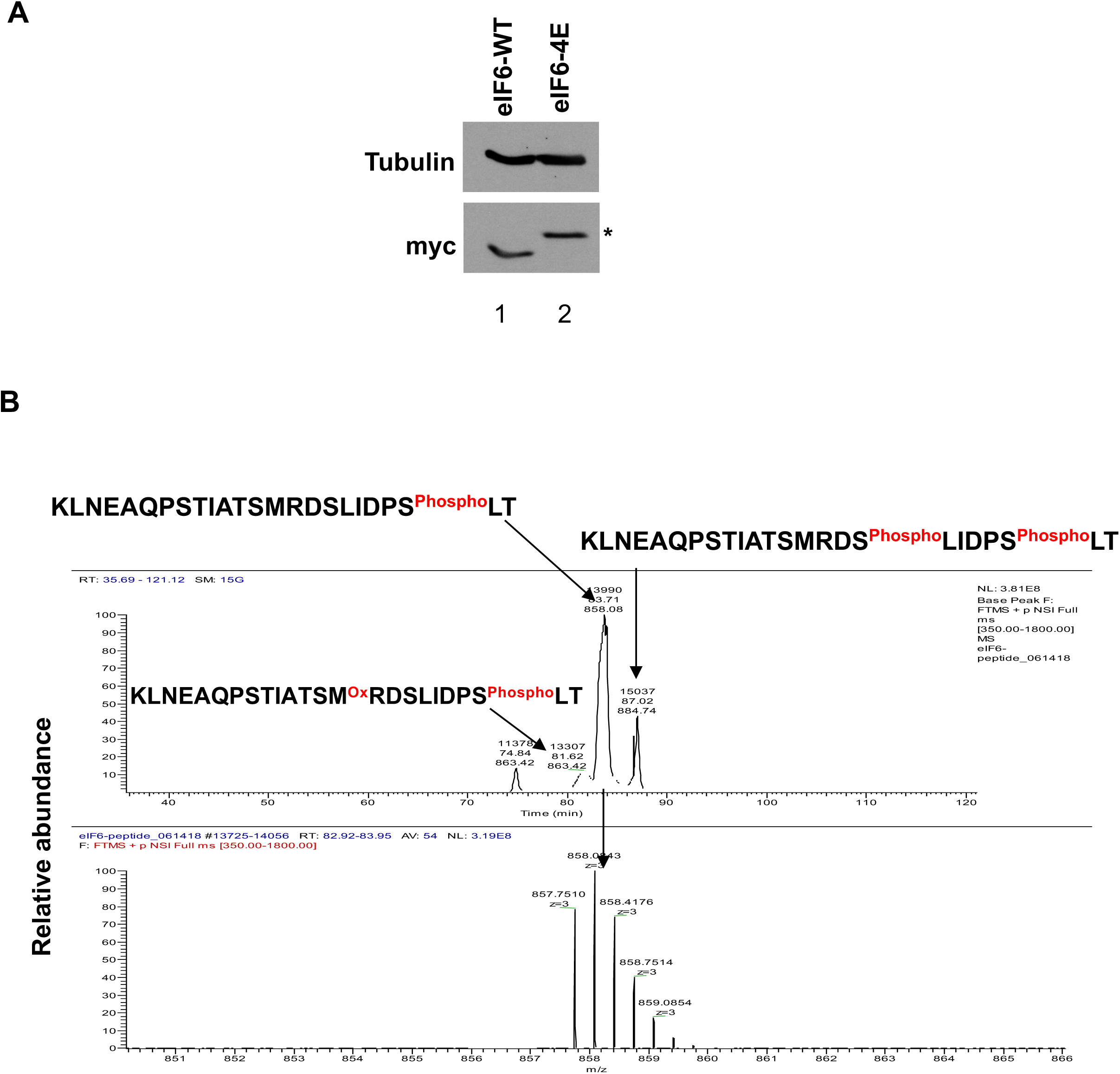
MS analysis of the synthetic phosphopeptide. **A.** HCT116 cells transfected with either myc-tagged eIF6-WT or phosphomimic-eIF6-4E mutant and whole cell lysates were subjected to western blotting using antibodies targeted against myc or anti-αTubulin (loading control). Asterix indicates the slower migrating species. **B.** MS analysis of the synthetic phosphopeptide was carried out as indicated in Fig. 2E. Extracted Ion Chromatograms and mass spectrum show multiple versions of the synthetic peptide and the relative abundance of the indicated synthetic peptides post *in vitro* phosphorylation.

**Figure S4.**
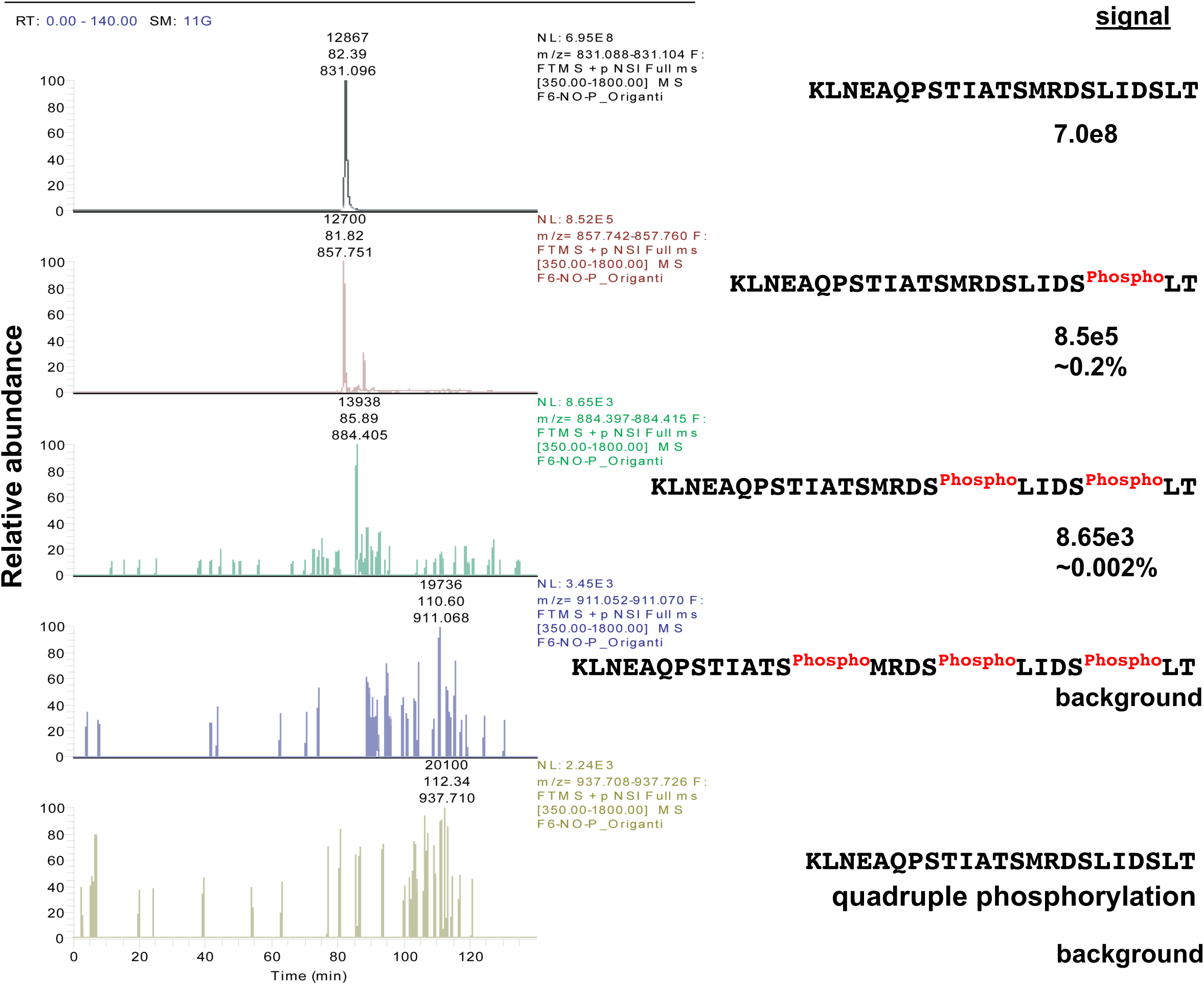
MS analysis of the synthetic non-primed peptide. For *nano*LC-MS/MS analysis, an eIF6 phosphopeptide (23 amino acids) carrying a phosphate lacking the priming phosphorylation was synthesized (Aapptec) and incubated with recombinant GSK3β. Extracted ion chromatograms show relative abundance levels of singly (pre-charged) and multiple residues phosphorylated by GSK3β in the region of interest.

**Figure S5.**
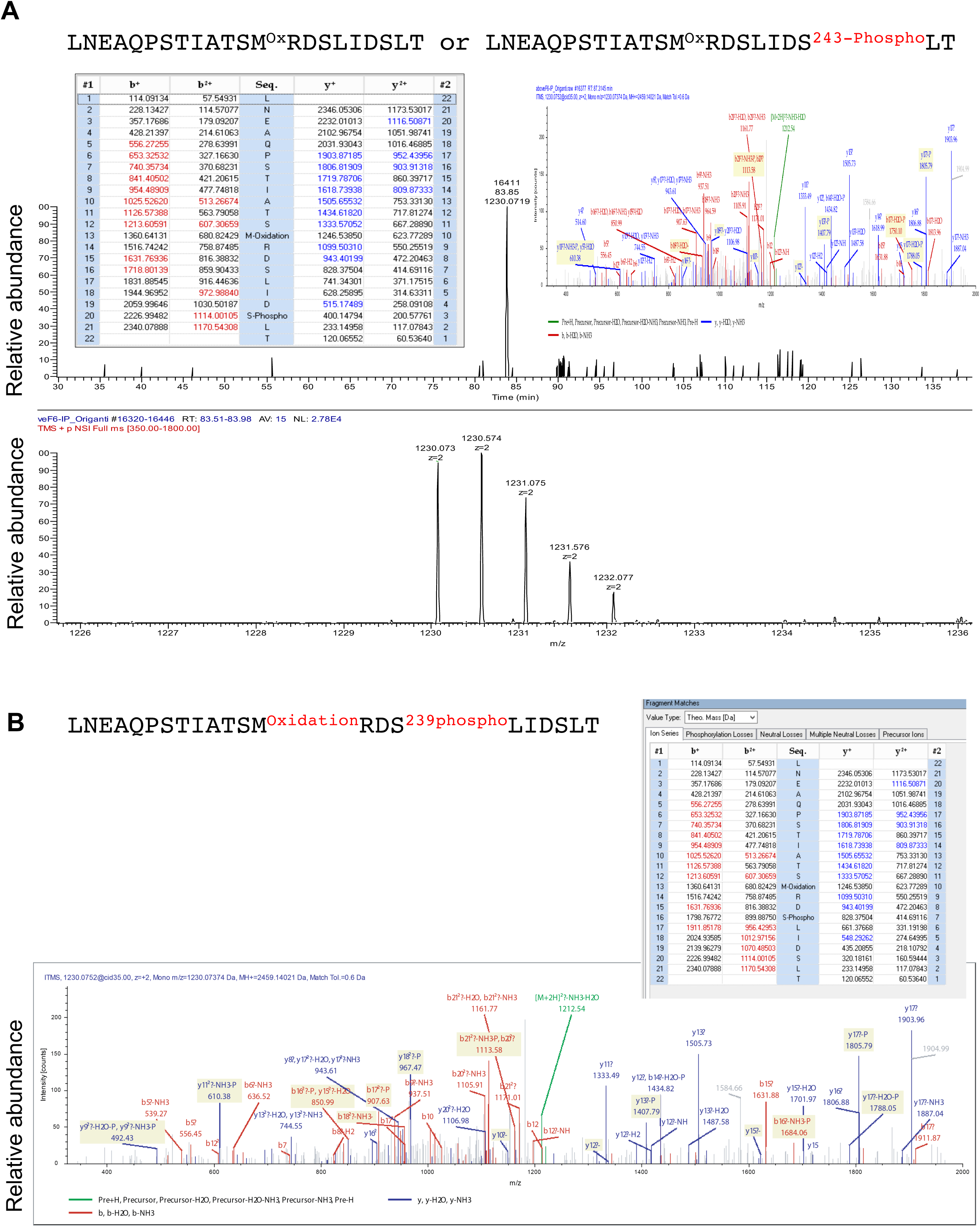
MS/MS analysis identified phosphorylation of the C-terminal tail of eIF6 in response to serum starvation *in vivo*. **A.** Mass Spectral (MS) analysis was carried out on myc-eIF6 immunoprecipitated from HCT116 cells that were serum starved for 24 hours. Samples were excised from coomassie-stained acrylamide gel and analyzed by *nano*LC-MS/MS analysis. Plot shows the EIC and MS/MS spectrum along with matched fragmentation mass table showing specific S243 phosphorylation in the region of interest. **B.** EIC and MS/MS fragmentation spectra shows *in vivo* phosphorylation of S239 site on myc-eIF6. Myc-eIF6 was immunoprecipiated from serum-starved HCT116 cells and subjected to MS analysis. Sequence indicates the C-terminal peptide showing S239 phosphorylation.

**Figure S6.**
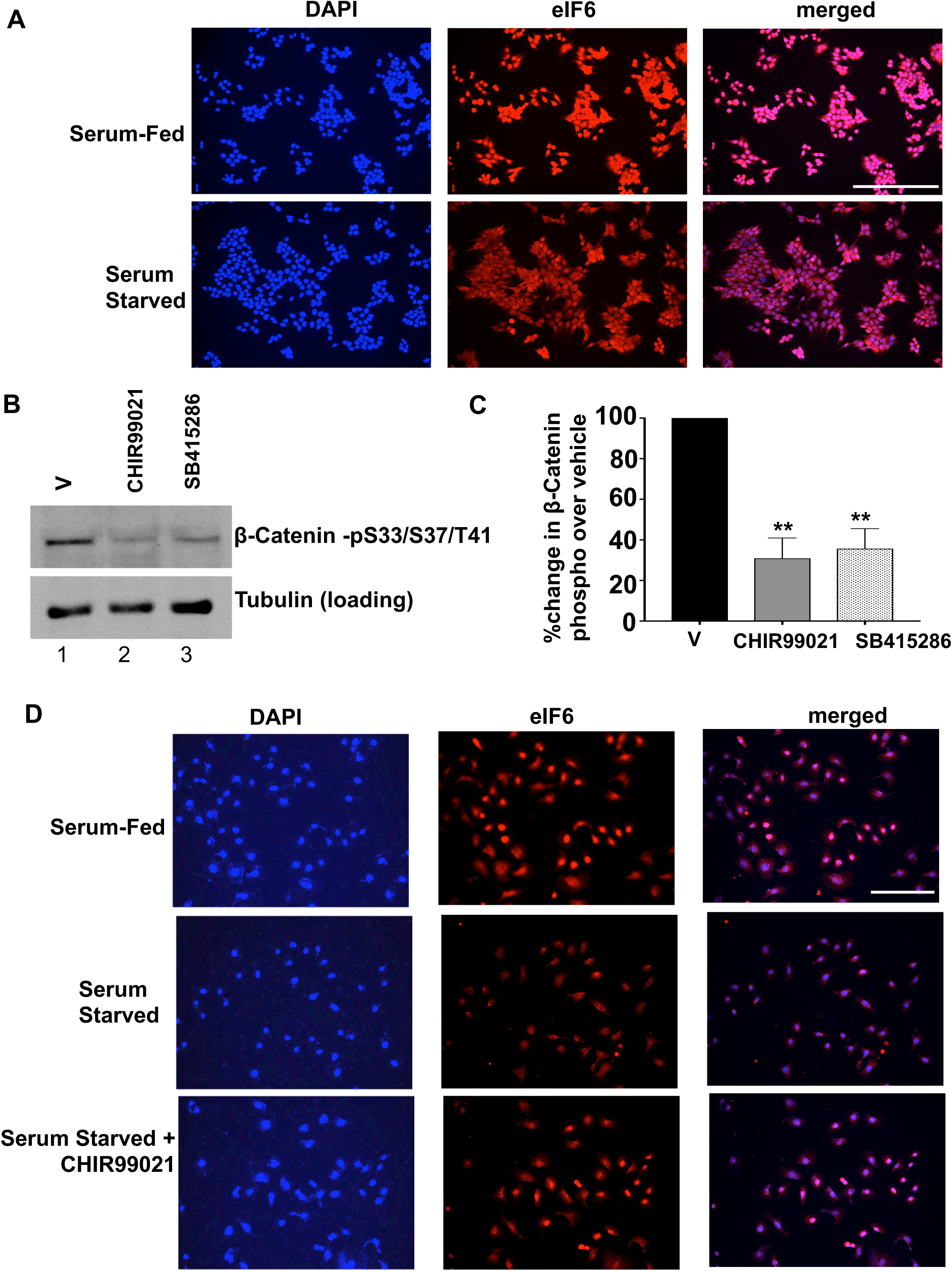
GSK3 regulates the cytoplasmic accumulation of eIF6 in response to serum starvation in normal RIE-1 cells. **A.** HCT116 cells were fed with 10% FBS or starved in 0.1% FBS for 24 hours. Cells were fixed and stained for endogenous eIF6 (red) using anti-eIF6 monoclonal antibody (Santa Cruz Biotechnology) and nuclei were stained with DAPI (blue) and analyzed by immunofluorescence microscopy (n=3). Scale bar = 250μm. **B.** HCT116 cells were serum-starved in 0.1% FBS for 24 hours and treated with either vehicle (0.1% DMSO) (lane 1), or 10μM CHIR99021 (lane 2), or 25μM SB415286 (lane 3) for 3hours. Whole cell lysates were then analyzed by western blotting using anti-β-Catenin-phospho-S33/S37/T41 antibody and anti-αTubulin antibody (loading control) (n=3). **C.** Blots represented in A. were quantitated and values represent the SEM of three independent experiments. Percent change in β-Catenin-phospho-S33/S37/T41 levels relative to the vehicle control was plotted. The differences were found to be significant as determined by an unpaired two-tailed t-Test with p=0.0023 for CHIR99021 versus vehicle and p=0.0029 for SB415286 versus vehicle. **D.** Normal RIE-1 cells were serum starved in 0.1% FBS for 24 hours and treated with either vehicle or 10μM CHIR99021 (4 hours). Cells fed with 10% FBS and treated with vehicle were used as the serum-fed controls. Cells were fixed and stained for endogenous eIF6 (red) and nuclei were stained with DAPI (blue) (n=3), and analyzed by immunofluorescence microscopy. Scale bar =100μm.

**Figure S7.**
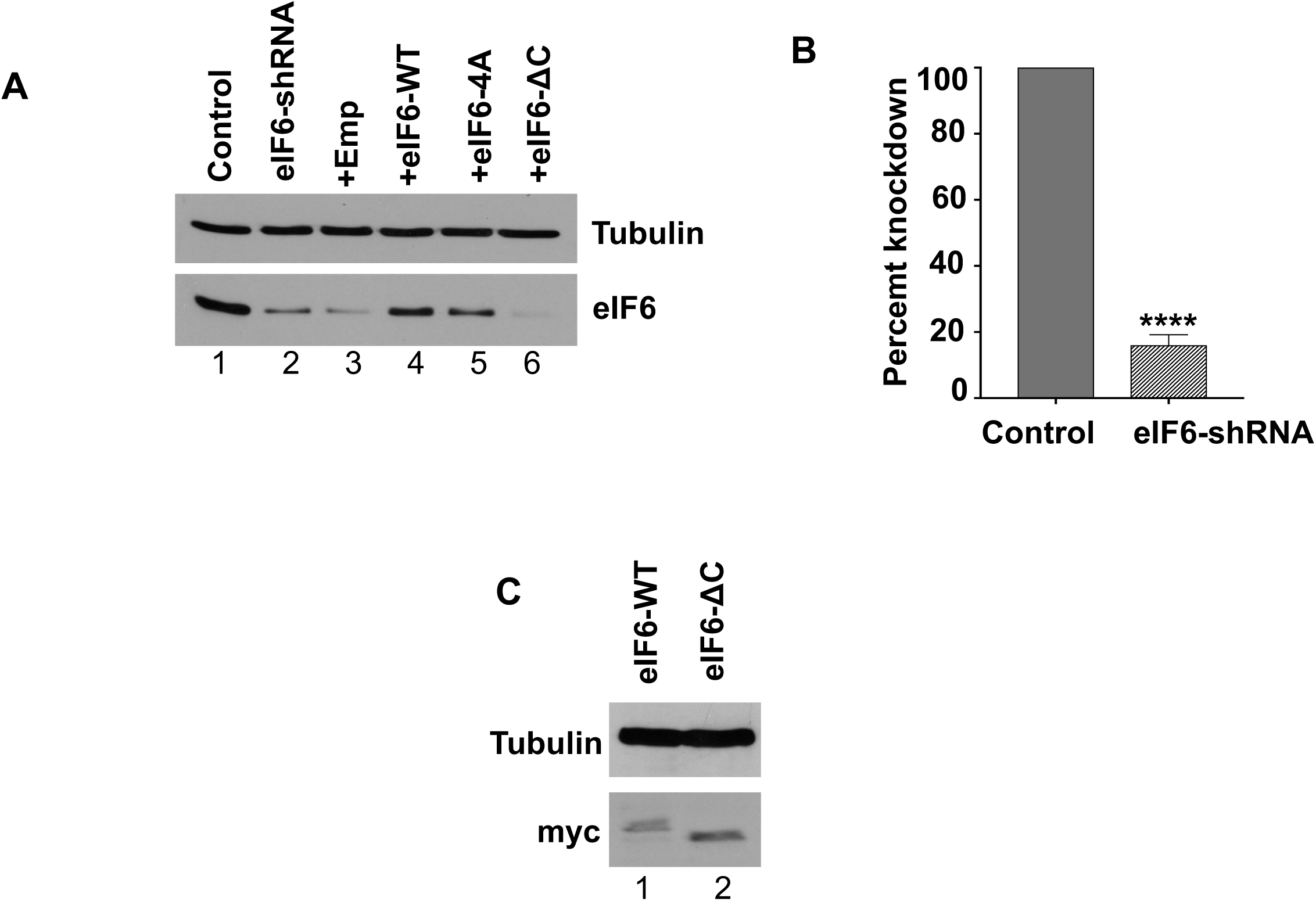
Generation of stable lines expressing eIF6-WT, eIF6-4A and eIF6-ΔC. **A.** Western blot of the HCT116 control, eIF6-KD (knocked down) cell line stably expressing eIF6 shRNA, and eIF6-KD line stably transfected with empty pcDNA3.1 vector, or pcDNA3.1 vector expressing myc-tagged eIF6-WT, eIF6-4A or eIF6-ΔC line, was probed with either anti-eIF6 monoclonal antibody or anti-Tubulin antibody (loading control). The anti-eIF6 monoclonal antibody (Cell Signaling) is targeted against the C-terminal tail of eIF6 and is unable to recognize the eIF6-ΔC mutant. (n=3) **B.** Blots represented in A. were quantitated and the percent knockdown of eIF6 relative to the untransfected control was plotted. Values represent the standard error of the mean (SEM) of three independent experiments and asterisks indicate a significant difference between eIF6-shRNA and the control with p <0.0001 as determined by an unpaired two-tailed t-Test. **C.** Myc-tagged eIF6-WT (lanes 1 and 2) or eIF6-ΔC mutant (lanes 3 and 4) were stably expressed in eIF6-KD cell line. Serum-fed eIF6-WT or eIF6-ΔC cells were analyzed by western blotting and probed with the indicated antibodies. (n=3)

**Figure S8.**
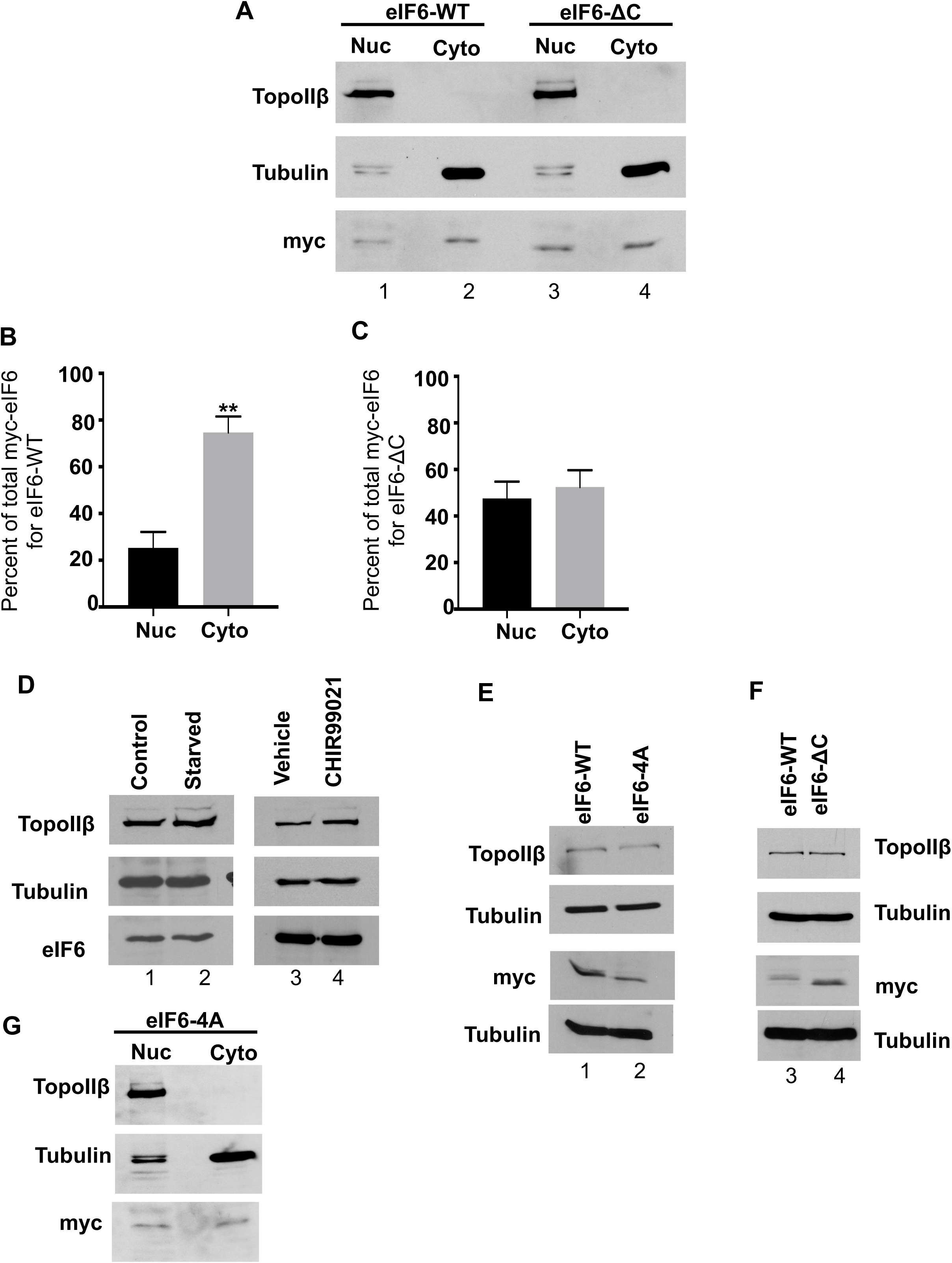
The eIF6-ΔC mutant does not show enhanced nuclear-cytoplasmic accumulation in response to serum starvation. **A.** Myc-tagged eIF6-WT or eIF6-ΔC mutant were stably expressed in the eIF6-KD cell line. Cells were serum starved for 24 hours followed by subcellular fractionation. Samples were analyzed by western blotting and blots were probed with anti-myc, anti-αTubulin, and anti-Topoisomerase IIβ antibodies. **B and C**. Blots represented in **A.** were quantitated and values represent the SEM of three independent experiments. Percent of nuclear and cytoplasmic fractions relative to the total (sum of nuclear and cytoplasmic eIF6 levels) are plotted. The difference in myc-eIF6 levels between the nuclear and cytoplasmic fractions for eIF6-WT (**B**) was found to be significant as determined by an unpaired two-tailed t-Test and asterisks indicate: p=0.007, but no significant difference was found for the eIF6-ΔC mutant (**C**). **D.** Whole cell lysates of samples subjected to nuclear-cytoplasmic fractionation in Fig. 5A (n=4) (lanes 1, 2) and Fig. 5D (n=3) (lanes 3, 4) were assayed for total levels of eIF6, αTubulin and Topoisomerase IIβ by western blotting. **E.** Whole cell lysates of samples subjected to nuclear-cytoplasmic fractionation in Fig. 5G were assayed for total levels of myc, αTubulin and Topoisomerase IIβ by western blotting (n=3). **F.** Whole cell lysates of the samples subjected to subcellular fractionation in Fig. S8C. were analyzed by western blotting and were probed with the indicated antibodies (n=3). **G.** Image represents the nuclear and cytoplasmic fractions of serum-fed HCT116 cells stably expressing myc-eIF6-4A. (n=3). Cytoplasmic and nuclear fractions were analyzed by western blotting using the indicated antibodies.

**Figure S9.**
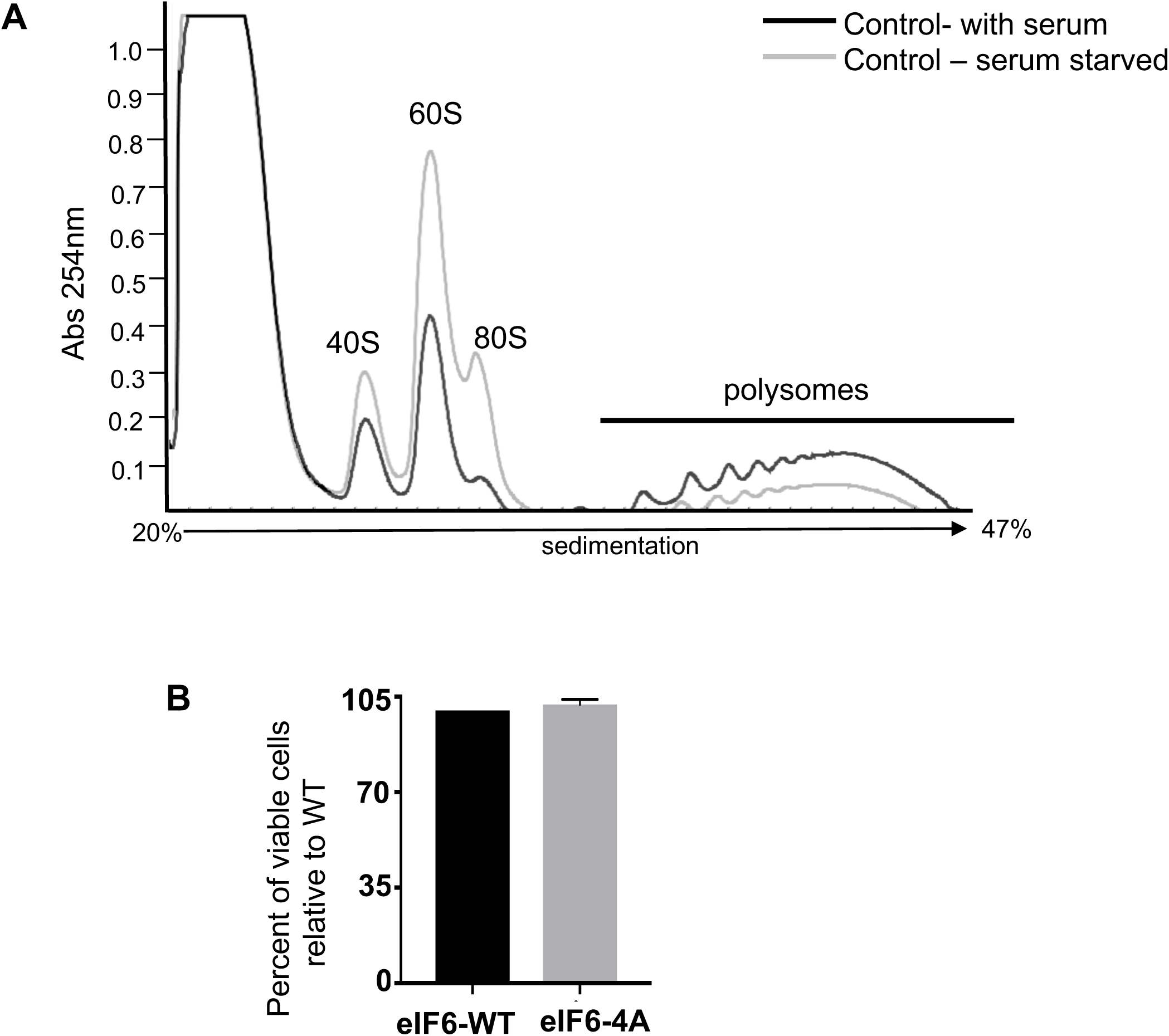
Polysome profile analysis of the eIF6-ΔC mutant compared to eIF6-WT under serum starved conditions. **A.** Representative polysome profile of HCT116 cells that were serum starved for 24 hours was overlaid with the polysome profile of serum-fed cells. Profiles are representative of three independent experiments. **B.** Bar graph represents the percentage of viable HCT116 cells stably expressing eIF6-4A mutant relative to eIF6-WT. Cells were cultured in the presence of serum and subjected to trypan blue dye exclusion assay. Graph represents the standard error of the mean (SEM) of three independent experiments. No significant difference was found between eIF6-WT and eIF6-4A mutant as determined by an unpaired two-tailed t-Test.

## Materials and Methods

### Cell culture and transfections

HCT116 cells (human colorectal carcinoma line) (America Type Culture Collection - ATCC) were maintained in McCoy’s 5A medium supplemented with 10% fetal bovine serum (FBS), and 100U/mL penicillin and 100μg/mL streptomycin (Gibco). HEK293T/17 and HeLa cells (ATCC) were maintained in Dulbecco’s Modified Eagle Medium supplemented with 10% FBS, and 100U/mL penicillin and 100μg/mL streptomycin (Gibco). For transfections, about 2 × 10^6^ HCT116 and 2.5 × 10^6^ 293T cells were plated per 60 mm dish. Each dish was transfected with 4μg of plasmid DNA and 20μL of lipofectamine 2000 (Invitrogen). 24 hours later, cells were washed twice in phosphate buffered saline (PBS), and serum starved for 4 hours using respective media with 0.1% FBS only. 4 hours later, cells were washed twice in PBS and collected in mammalian cell lysis buffer (MCLB) (50mM Tris-Cl pH 8.0, 5mM ethylenediaminetetraacetic acid (EDTA), 0.5% IGEPAL, 150mM sodium chloride) that was supplemented with the following inhibitors just before lysis: 1mM phenylmethylsufonyl fluoride (PMSF), 1mM sodium fluoride, 10mM β-glycerophosphate, 1mM sodium vanadate, 2mM Dithiothreitol (DTT), 1X protease inhibitor cocktail (Sigma Aldrich), 1X phosphatase inhibitor cocktail (Santa Cruz Biotechnology). Lysates were rocked for 15 minutes at 4°C, followed by centrifugation at 14,000 rpm for 10 minutes at 4°C.

### Plasmid constructions

To clone human eIF6, mRNA was extracted from MCF7 (ATCC) cells using RNAaqueous Total RNA Isolation kit (Invitrogen) and cDNA was synthesized using the Superscript First Strand cDNA synthesis kit (Invitrogen). Using polymerase chain reaction (PCR), eIF6 ORF was cloned into the pCMV-myc plasmid (Clontech) using the respective forward and reverse primers carrying EcoRI and XhoI restriction sites: 5’-TATAAGAATTCTAATGGCGGTCCGAGCTTCGTTCGAGAAC-3’, 5’-TATAACTCGAGTCAGGTGAGGCTGTCAATGAGGGAATC-3. GST-tagged GSK3β constructs were cloned into the pDEST™27 vector (ThermoFisher Scientific) as described previously^66^. All the pCMV and pcDNA3.1-myc-eIF6 phospho-site mutants were generated using the Q5^®^ Site-Directed Mutagenesis Kit (New England Biolabs) using the forward and reverse primers listed in Appendix Table S1. All plasmid constructs used in this study were verified by Sanger sequencing (Genewiz).

### Generation of stable cell lines expressing WT and mutant eIF6

HCT116 cells were transfected with pLKO.1 vector expressing either an eIF6 targeted shRNA (CCGGGTGCATCCCAAGACTTCAATTCTCGAGAATTGAAGTCTTGGGATGCACTTTTTG, Sigma - TRC327700) or a control non-targeting shRNA (Sigma) as described above. Stably transfected clones were selected in 2μg/mL puromycin and maintained in 0.5μg/mL puromycin. ∼30 clones were screened for eIF6 knockdown (eIF6-KD) by western blotting. To express eIF6 WT and mutants in the eIF6-KD cell line, cells were stably transfected with pcDNA3.1 plasmid expressing either the myc-tagged eIF6-WT, eIF6-4A, eIF6-ΔC or the empty vector. To evade shRNA targeting, mismatches were generated in the myc-eIF6-WT, myc-eIF6-4A or myc-eIF6-ΔC plasmids using site directed mutagenesis. All stably transfected clones were selected in 500μg/ml Geneticin and 0.5μg/mL puromycin. ∼25 clones were screened for myc-eIF6-WT or mutant expression by western blotting. All stable lines were maintained in 100μg/ml Geneticin and 0.5μg/mL puromycin.

### Cell viability assays

7,000 HCT116 cells expressing eIF6-WT or eIF6-4A were cultured per well of a 96-well plate in McCoy’s culture media without phenol red. After overnight incubation, cells were serum starved for 24 hours in McCoy’s media supplemented with 0.1% FBS. 20μL of MTS solution (CellTiter 96^®^ Aqueous One Solution Reagent, Promega) was added to each well and plates were incubated for 1 hour. Absorbances were read at 490 nm (Spectra Max i3x, Molecular Devices). To obtain background-corrected absorbances, average absorbance values of the control wells containing media and MTS only were subtracted from all other absorbances. For trypan blue assay, HCT116 cells expressing eIF6-WT or eIF6-4A were plated as indicated for polysome profile assays. Cells suspended in culture media were mixed with trypan blue dye at 1:1 ratio and the number of blue-stained (dead) cells and unstained (viable) cells were counted using a hemocytometer.

### Immunoprecipitation

For immunoprecipitation of myc-eIF6, or myc-empty vector (Fig. 1B), lysates with 1.2 mg of total protein were suspended in a final volume of 500μL of MCLB buffer supplemented with inhibitors. Lysates were pre-cleared with 20μL of protein A/G plus-conjugated agarose beads (Santa Cruz Biotechnology) and 1μL of normal mouse IgG (Santa Cruz Biotechnology). Protein A/G plus-agarose beads were washed in MCLB buffer before use. For pre-clearing, lysates were rotated for 20 minutes at 4°C. Pre-cleared lysates were collected by centrifugation at 3500 rpm for 3 minutes at 4°C, and the beads were discarded. For immunoprecipitation, pre-cleared samples were incubated with 20μL of protein A/G plus-agarose beads and 11μL of myc-tag antibody (9E10) (Santa Cruz Biotechnology) and rotated overnight at 4°C. Samples were washed three times in MCLB buffer and the beads were collected by centrifugation at 3000 rpm for 3 minutes and washed for a fourth time in incomplete kinase (IK) buffer (50mM Tris, pH7.4, 10mM MgCl_2_, 2mM DTT), and after the final wash, beads were suspended in 45μL of IK buffer. Immunoprecipitation of the myc-tagged phospho-site mutants of eIF6, eIF6-CΔ20, and myc-eIF6-WT was carried out as described above except that 1.5mg of total protein was used for the assay. For the myc-eIF6-ΔC36 mutant, owing to a small reduction in its expression, 2mg of total protein was used for the assay. This ensured that the total eIF6 protein input for the kinase assays were comparable between eIF6-WT and the eIF6-ΔC36 mutant.

### *In vitro* kinase assay

For the *in vitro* kinase assays, myc-tagged versions of eIF6 were immunoprecipitated as indicated above and the immunoprecipitated beads were incubated with, or without 350 units of recombinant GSK3β (rabbit skeletal muscle) (New England BioLabs) and 1X hot kinase buffer (50mM Tris, pH 7.4, 50mM MgCl_2_, 10mM DTT, 50μM cold ATP, γ-^32^P-ATP-1μCi), and incubated at 30°C for 30 minutes. Kinase reactions were also set-up for the analysis of phospho-site mutants of eIF6 and eIF6-WT control as detailed above except that the reactions were incubated at 30°C for 25 minutes to account for time-dependent kinetic differences. Kinase reactions were analyzed by SDS-PAGE and exposed to X-ray film for autoradiography. For western blotting of the kinase assay, duplicate kinase reactions were simultaneously set-up as described above except that the reactions were set-up using the cold kinase buffer (50mM Tris, pH 7.4, 50mM MgCl_2_, 10mM DTT, 250μM cold ATP). Blots were probed with anti-eIF6 monoclonal antibody (1:1000), or anti-myc-tag monoclonal antibody (1:750), and anti-GSK3β monoclonal antibody (1:1000) (Cell Signaling Technology) diluted in Tris-buffered saline with 0.1% Tween-20 (TBS-T).

### Pull-down assay

To determine interactions with the endogenous eIF6, HCT116 cells were transfected with GST-GSK3β or empty vector as previously described. Cell lysis was carried out in MCLB buffer as described above and a small fraction was saved for analysis of input levels. For the pull-down assays, 1.1mg of total protein suspended in 500μL of MCLB buffer and supplemented with inhibitors, was incubated with 50μL of glutathione agarose resin beads (Gold Biotechnology) and rotated overnight at 4°C. To determine interactions with the GST-only control, recombinant GST was purified from BL21 Codon-Plus *E. coli* using glutathione affinity chromatography, and 500ng of recombinant GST was incubated with lysates transfected with the empty vector control at 4°C. Next day, samples were washed three times in MCLB buffer supplemented with inhibitors and resuspended in (30μL) of MCLB buffer and 2X Laemmli buffer. Samples were then analyzed by western blotting. For the pull-down of GST-GSK3β-WT and GST-GSK3β-R96A, 293T cells were transfected with the respective vectors and 1.9mg of total protein was used for the pull-down analysis. Pull-downs were carried out as described above except that the active kinase was eluted from the glutathione agarose beads using glutathione (10mM reduced glutathione in 50mM Tris-HCl, pH 8.0). For the kinase assay, fresh eluates were incubated with myc-eIF6 that was immunoprecipitated (2mg total protein) from 293T cells.

### Western blotting

For western blotting, proteins were resolved by SDS-polyacrylamide gel electrophoresis (SDS/PAGE). For all blots, about 20-40μg of total protein was loaded per lane, except for the kinase reactions where samples were prepared as described above. Proteins were transferred onto nitrocellulose membranes (0.45 micron, Bio-Rad Laboratories). Blots were blocked either for 1 hour or overnight in 5% non-fat dry milk diluted in TBS-T. The following primary antibodies dissolved in TBS-T were used: anti-GST antibody (1:1000, overnight), anti-αTubulin antibody (1:1000, overnight), anti-β-actin antibody (1:3000, 1hr), anti-eIF6 antibody (1:1000, overnight), anti-GSK3β antibody (1:1000, overnight) anti-phospho GSK3β (Ser-9) antibodies (1:1000, overnight), anti-myc-tag antibody (1:1000, overnight) (Cell Signaling Technology). To assay for GSK3β interaction with endogenous eIF6, blots were probed with anti-eIF6 monoclonal antibody (1:500) and incubated overnight (Cell Signaling Technology). To assay for β-catenin phosphorylation, 100μg of total protein was loaded per lane and blots were probed with anti-phospho-β-catenin (Ser33/37/Thr41) antibody (1:750 in 5% milk-TBS-T) and incubated overnight. Secondary antibodies (Jackson ImmunoResearch) were diluted 1:30000 in TBS-T. Blots were developed using ECL western blotting substrate (Pierce). For western blotting of nuclear and cytoplasmic fractions, 20μg of total protein was loaded per lane and probed with the following primary antibodies dissolved in TBS-T: Topoisomerase IIβ (1:1000, overnight) (BD Biosciences), αTubulin (DMIA) (1:3000, 1hr) and anti-eIF6 antibody (D1696) (1:1000, overnight) (Cell Signaling Technology).

### Mass Spectrometry

For mass spectrometry, 7 × 10^6^ 293T cells were plated per 10cm plate and transfected as indicated before. 24 hours later, cells were serum starved in DMEM containing 0.1% FBS for 3 hours. Cells were lysed in MCLB buffer supplemented with inhibitors. Immunoprecipitations were carried out as previously described using 3mg of total protein. For the kinase reactions, 15μL of myc-eIF6-bound beads were incubated with/without recombinant GSK3β (NEB) and suspended in cold kinase buffer and incubated at 30°C for 30 minutes. Reactions were washed twice in incomplete kinase buffer, and resuspended in incomplete kinase buffer and 1.5X Laemmli buffer. Samples were boiled and analyzed by SDS-PAGE followed by Coomassie staining. Stained gels were sent to the University of Wisconsin-Madison Mass Spectrometry facility.

#### Enzymatic “In Gel” Digestion

“In gel” digestion and mass spectrometric analysis was done at the Mass Spectrometry Facility [Biotechnology Center, University of Wisconsin-Madison]. In short, Colloidal Coomassie G-250 stained gel pieces were de-stained twice for 5 minutes in MeOH:H_2_0:NH_4_HCO_3_ [50%:50%:100mM], dehydrated for 5 minutes in ACN:H_2_0:NH_4_HCO_3_ [50%:50%:25mM] then once more for 30 seconds in 100% ACN, dried in a Speed-Vac for 2minutes, rehydrated completely and reduced in 25mM DTT (in 25mM NH_4_HCO_3_) for 30 minutes at 56°C, alkylated by solution exchange with 55mM Iodoacetamide (in 25mM NH_4_HCO_3_) in the dark at room temperature for 30 minutes, washed once in 25mM NH_4_HCO_3_, dehydrated twice for 5 minutes in ACN:H_2_0:NH_4_HCO_3_ [50%:50%:25mM] then once more for 30 seconds in 100% ACN, dried in a Speed-Vac again and finally rehydrated with 20μL of trypsin solution [10ng/μL trypsin (PROMEGA) in 25mM NH_4_HCO_3_ and 0.01% ProteaseMAX w/v (PROMEGA)]. Additional 30μL of digestion solution [25mM NH_4_HCO_3_ and 0.01% ProteaseMAX w/v (PROMEGA)] was added to facilitate complete rehydration with excess overlay needed for peptide extraction. The digestion was conducted for 3 hours at 42°C. Peptides generated from digestion were transferred to a new tube and acidified with 2.5% Triflouroacetiacid (TFA) to 0.3% final concentration. Gel pieces were extracted further with ACN:H_2_O:TFA [70%:29.25%:0.75%] for 10 minutes and solutions combined then dried completely in a Speed-Vac (∼15 minutes). Extracted peptides were solubilized in 30µL of 0.1% formic acid and degraded in ProteaseMAX and removed via centrifugation [max speed, 10 minutes]. 10µL was taken for immediate solid phase extraction (ZipTip^®^ C18 pipette tips Millipore) according to manufacturer protocol. Remaining 20µLwas subjected to a secondary digestion with endoproteinase AspN (Roche) where pH was adjusted with 2.5µL 500mM NH_4_HCO_3_ and 1µL of 40ng/µL AspN stock added. Digestion was carried out for 2 hours at 37°C subsequently acidified with 2.5% TFA to 0.3% final and solid phase extracted as conventional tryptic digest. Peptides eluted off ZipTip C18 tips with ACN:H_2_O:TFA [70%:29.9%:0.1%] were dried and finally solubilized in 8µL of 0.1% formic acid.

#### NanoLC-MS/MS

Peptides were analyzed by nano liquid chromatography tandem mass spectrometry using the Agilent 1100 nanoflow system (Agilent) connected to a hybrid linear ion trap-orbitrap mass spectrometer (LTQ-Orbitrap Elite™, Thermo Fisher Scientific) equipped with an EASY-Spray™ electrospray source. Chromatography of peptides prior to mass spectral analysis was accomplished using capillary emitter column (PepMap® C18, 3µM, 100Å, 150×0.075mm, Thermo Fisher Scientific) onto which 3µL of extracted peptides was automatically loaded. NanoHPLC system delivered solvents A: 0.1% (v/v) formic acid, and B: 99.9% (v/v) acetonitrile, 0.1% (v/v) formic acid at 0.50 µL/min to load the peptides (over a 30 minute period) and 0.3µl/min to elute peptides directly into the nano-electrospray with gradual gradient from 3% (v/v) B to 20% (v/v) B over 17 minutes and concluded with 5 minute fast gradient from 20% (v/v) B to 50% (v/v) B at which time a 4 minute flush-out from 50-95% (v/v) B took place. As peptides eluted from the HPLC-column/electrospray source survey MS scans were acquired in the Orbitrap with a resolution of 120,000 followed by MS2 fragmentation of 20 most intense peptides detected in the MS1 scan from 350 to 1800 m/z; redundancy was limited by dynamic exclusion.

#### Data analysis

Raw MS/MS data was converted to mgf file format using MSConvert (ProteoWizard: Open Source Software for Rapid Proteomics Tools Development). Resulting mgf files were used to search against Uniprot Human amino acid sequence database containing a list of common contaminants and decoy entries (134,183 total entries) using in-house *Mascot* search engine 2.2.07 (Matrix Science) with variable Serine and Threonine phosphorylation, Methionine oxidation, Asparagine and Glutamine deamidation plus fixed Cysteine carbamidomethylation. Peptide mass tolerance was set at 15 ppm and fragment mass at 0.6 Da.

#### Mass spectrometry of phosphopeptide and non-phosphorylated peptide

100µM of the synthesized phosphopeptides (Aapptec) or non-phosphorylated peptides were incubated with 350units of recombinant GSK3β (New England Biolabs) and cold kinase buffer. Reactions were incubated for 20 minutes at 30°C. Reactions were then quenched by 40mM EDTA and by freezing.

#### NanoLC-MS/MS

100µM of synthesized version of human eIF6 phosphopeptide was acidified with 2.5% TFA to 0.4% final and solid phase extracted (ZipTip C18 pipette tips, Millipore) according to manufacturer protocol. Peptide was eluted off ZipTip C18 tip with 2µl of ACN:H_2_O:TFA [70%:29.9%:0.1%] and diluted to 20µL final volume with 0.1% formic acid. Mass spectrometric analysis followed using the Agilent 1100 nanoflow system (Agilent) connected to a hybrid linear ion trap-orbitrap mass spectrometer (LTQ-Orbitrap Elite™, Thermo Fisher Scientific) equipped with an EASY-Spray™ electrospray source. Chromatography of peptides prior to mass spectral analysis was accomplished using capillary emitter column (PepMap C18, 3µM, 100Å, 150×0.075mm, Thermo Fisher Scientific) onto which 2µL of extracted peptides was automatically loaded. NanoHPLC system delivered solvents A: 0.1% (v/v) formic acid, and B: 99.9% (v/v) acetonitrile, 0.1% (v/v) formic acid at 0.50 µL/min to load the peptides (over a 30 minute period) and 0.3µL/min to elute peptides directly into the nano-electrospray with gradual gradient from 3% (v/v) B to 20% (v/v) B over 17 minutes and concluded with 5 minute fast gradient from 20% (v/v) B to 50% (v/v) B at which time a 4 minute flush-out from 50-95% (v/v) B took place. As peptides eluted from the HPLC-column/electrospray source survey MS scans were acquired in the Orbitrap with a resolution of 120,000 followed by MS2 fragmentation of 20 most intense peptides detected in the MS1 scan from 350 to 1800 m/z; redundancy was limited by dynamic exclusion. Raw MS/MS data was converted to mgf file format using MS Convert (ProteoWizard: Open Source Software for Rapid Proteomics Tools Development). Resulting mgf files were used to search specifically against human eIF6 sequence using in-house *Mascot* search engine 2.2.07 (Matrix Science) with variable Serine and Threonine phosphorylation. Peptide mass tolerance was set at 15 ppm and fragment mass at 0.6 Da. Dynamic phosphorylation cascade distributions were manually interrogated for precursor ion abundance and quality of MS/MS fragmentation.

#### Mass spectrometry of serum starved samples

For *in vivo* mass spectrometry analysis, myc-eIF6 was immunoprecipitated from HCT116 cells that were serum starved for 24 hours. Immunoprecipitations were carried out as previously described except that 6mg of total protein was used for analysis, and each band of a doublet of myc-eIF6 including a slower migrating species was excised from coomassie-stained gel and subjected to MS analysis as described above. Phosphopeptides were only detected in the slower migrating/super shifted band.

#### Immunofluorescence staining and microscopy

50,000 HCT116 cells were plated in each well of a 12-well plate carrying poly-D-lysine-coated glass coverslips (Neuvitro). After overnight incubation, cells were either serum-fed with McCoy’s 5A medium containing 10% FBS or serum starved in media containing 0.1% FBS, and 24 hours later, treated with either 0.1% dimethyl sulfoxide (DMSO) (vehicle), or 10µM CHIR99021 (Tocris), or 25µM SB415286 (Tocris). Cells were fixed overnight in 2% paraformaldehyde/PBS. For immunostaining, fixed cells were permeabilized with 2% Triton X-100/PBS for 20 minutes and incubated in blocking buffer (2% BSA/0.1% IGEPAL/PBS) for 30 minutes. The following primary antibodies diluted in the blocking buffer were used for staining: anti-eIF6 monoclonal antibody (D16E9) (1:300) (Cell Signaling Technology), or anti-eIF6 monoclonal antibody (1:350) (Santa Cruz Biotechnology) and incubated in a humidified chamber at room temperature (RT) for 1 hour. Coverslips were washed with 0.1% IGEPAL/PBS and incubated with Cy3-conjugated affinipure donkey anti-rabbit IgG secondary antibody (Jackson ImmunoResearch) diluted in blocking buffer (1:300) at RT for 1 hour in the dark. Coverslips were washed in 0.1% IGEPAL/PBS and mounted using ProLong Gold antifade reagent with 4’-6-Diamidino-2-phenylindole (DAPI) (Invitrogen). Immunofluorescence studies in HeLa cells were carried out as described above. Staining of normal rat intestinal epithelial cells (RIE-1) was carried out as described above except that 30,000 cells were plated per well, and cells were probed with anti-eIF6 monoclonal antibody (D16E9) (1:200) (Cell Signaling Technology). RIE-1 cells were cultured and maintained as described previously^67,68^. For imaging, slides were analyzed using the Leica DM6 B upright fluorescent microscope. Images were acquired using Leica DFC 3000G (Bin 1×1, Gamma1) camera and processed by Leica LAS X imaging software.

#### Subcellular fractionation

700,000 HCT116 cells were plated per 100mm dish and 24 hours later, cells were serum starved in media containing 0.1% FBS or fed with media containing 10% FBS for an additional 24 hours. To assay the effect of inhibitors, serum starved cells were treated with either 0.1% DMSO (vehicle) or 10µM CHIR99021 (Tocris) for 5 hours. For subcellular fractionation of WT and mutants, 1.2 million cells were plated per 60mm dish. To assay for total protein input, cells were lysed in MCLB buffer supplemented with inhibitors as previously described. To extract cytoplasmic fractions, cells were collected in cell lysis buffer (10 mM HEPES, pH 7.5, 10 mM KCl, 0.1 mM EDTA, 0.5% IGEPAL) supplemented with the following inhibitors: 10mM β-glycerophosphate, 1mM sodium vanadate, 2mM DTT, 1X protease inhibitor cocktail (Sigma Aldrich), 1X phosphatase inhibitor cocktail (Santa Cruz Biotechnology) and 0.5mM PMSF. Nuclear-cytoplasmic fractionation was carried out as described before^69^. For cytoplasmic fractionation, lysates were incubated on ice for 15 minutes with intermittent gentle mixing and vortexed on high for 10 seconds at 14000 rpm. Cells were centrifuged at 12,000 x g for 10 minutes at 4°C and cytoplasmic lysates were collected. For nuclear extraction, nuclear pellets were washed four times in cell lysis buffer and centrifuged at 3000 rpm for 5 minutes at 4°C. Nuclear pellets were suspended in 75μL of nuclear extract buffer (20mM HEPES, pH 7.5, 400mM sodium chloride, 1mM EDTA,) supplemented with the following inhibitors: 10mM β-glycerophosphate, 1mM sodium vanadate, 2mM DTT, 1X protease inhibitor cocktail (Sigma Aldrich), 1X phosphatase inhibitor cocktail (Santa Cruz Biotechnology) and 0.5mM PMSF. Nuclear pellets were solubilized on ice for 30 minutes followed by centrifugation at 12,000 x g for 15 minutes at 4°C. Both nuclear and cytoplasmic fractions were analyzed by western blotting.

#### Polysome profile analysis

Polysome profile analysis was carried out as described previously except 1.5 million cells were plated per 10cm plate for eIF6-WT and eIF6-4A mutant serum-fed controls, and 2 million cells were plated per 10cm plate for serum starved samples (50% to 60% cell density)^67,68^. Cells were treated with 100µg/mL cycloheximide (Tocris) for 5 minutes at 37°C and washed with ice-cold 1X PBS buffer containing 100µg/mL cycloheximide on ice. Cells were lysed in the polysome lysis buffer (50mM HEPES pH 7.4, KCl-250mM, 5mM MgCl_2_, 250mM sucrose, 1% TritonX-100, 1% sodium deoxycholate, 100ug/mL cycloheximide, 100 units/mL Superasin (Invitrogen), 0.25X protease inhibitor cocktail EDTA free (Sigma) and 1X phosphatase inhibitor cocktail (Santa Cruz Biotechnology). About 10.5 OD A_260_ were loaded per linear sucrose gradient (20% to 47%). For each experiment, the same OD amount of lysate (600µL) was layered per sucrose gradient (20mM HEPES pH7.4, 200mM KCl, 10mM MgCl2, 100µg/mL cycloheximide, Rnase-free sucrose) and gradients were centrifuged at 35,000 rpm for 3 hours 20 minutes using a SW41Ti rotor (Beckman Coulter). Absorbance was followed at 254 nm. The Area under a Curve (AUC) was measured using the Peak Chart software (Brandel).

---

**Table S1.**
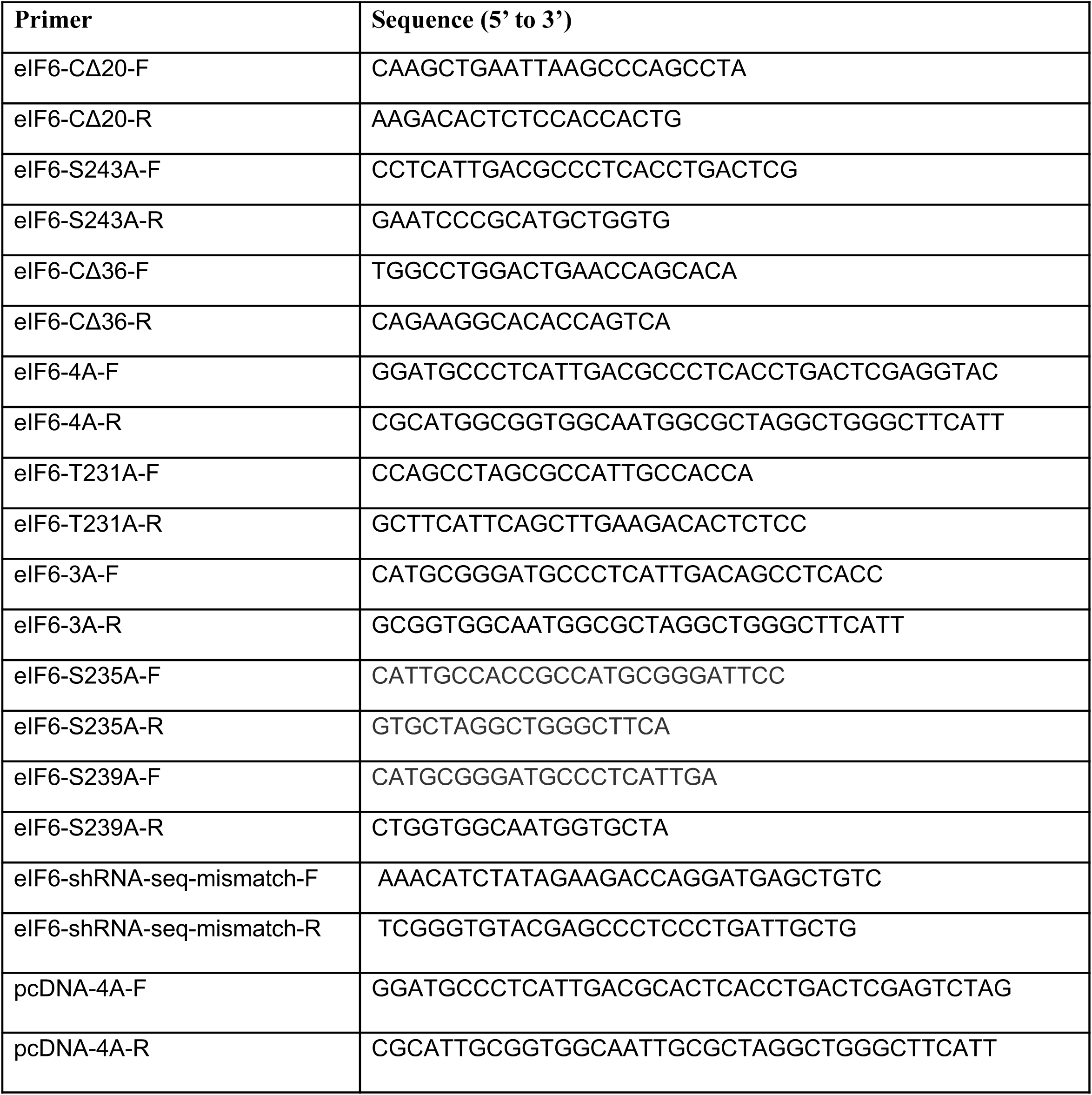
List of all forward (F) and reverse (R) primer names and sequences used in this study. eIF6-shRNA-seq-mismatch primers were used to generate mismatches in the eIF6-shRNA target recognition sequence in eIF6-WT and eIF6-mutants so that eIF6 expression could be rescued in HCT116 cells stably expressing eIF6-shRNA.

